# α2,3-sialylation on human naïve T cells restrains bispecific engager-mediated anti-tumor immunity

**DOI:** 10.64898/2026.08.02.742298

**Authors:** Johanna Nimmerfroh, Dinah Heiligensetzer, Michael Sandholzer, Anastasiya Börsch, Andreas Zingg, Christoph Schultheiss, Mascha Binder, Rafael Martinez Carrasco, Pablo Argüeso, Alfred Zippelius, Luana Guerra, Heinz Läubli

**Author notes:** Corresponding author: Heinz Läubli, Petersgraben 4, 4031 Basel, Switzerland.

## Abstract

Aberrantly elevated cell-surface sialylation, or hypersialylation, is a common feature of human cancers and contributes to immune evasion. Sialidase-based therapies have therefore emerged as a strategy to disrupt this glyco-checkpoint. Although the immunosuppressive role of tumor-associated sialylation is well established, how sialylation on human T cells shapes anti-tumor responses remains poorly defined. Here, we identify surface sialoglycans on T cells, particularly α2,3-linked structures, as a cell-intrinsic restraint on human T cell activation, proliferation, and effector function. *In vitro*, enzymatic desialylation enhanced T cell activation, proliferation, cytokine production, and bispecific T cell engager (TCE)-mediated tumor-cell killing in healthy donor PBMC co-cultures. In *ex vivo* cultures of primary chronic lymphocytic leukemia (CLL) PBMCs, sialidase treatment combined with the CD20-directed TCE glofitamab enhanced cytotoxic effector transcriptional programming in autologous T cells. Single-cell RNA sequencing combined with lectin-based CITE-seq linked treatment-induced transcriptional states to lectin-defined cell-surface glycan signatures within the same single-cell dataset. This integrated analysis revealed that naïve and, to a lesser extent, central memory T cells combined elevated baseline α2,3-sialylation signatures with the clearest transcriptional responses to glofitamab plus sialidase treatment. CD43 emerged as a major carrier of α2,3-linked sialoglycans, and its deletion attenuated sialidase-enhanced T cell activation. Together, these findings identify sialylation of the T cell surface as a subset-specific restraint on human TCE responses and provide a rationale for testing sialidase-TCE combinations designed to engage less-differentiated T cell populations.

**One-sentence summary:** Desialylation enhances bispecific T cell engager responses by relieving a sialoglycan-dependent restraint in human T cells.

## Introduction

The surface of mammalian cells is coated by a dense layer of glycans attached to proteins and lipids^1,2^. Many of these glycans terminate in sialic acids, forming sialoglycans that regulate cell-cell interactions and immune recognition^3^. In cancer, tumor-cell hypersialylation is a feature of malignant transformation and is associated with progression, metastasis, and immune evasion^4,5,6^. Tumor-cell sialoglycans can thereby act as glyco-checkpoints and therapeutic targets^7,8^, motivating enzyme-based strategies that remove terminal sialic acid residues from cell-surface glyco-conjugates. Sialidases have been delivered to tumors using engineered bacteria^9^, chimeric antigen receptor (CAR) T cells^10,11^, and antibody-derived constructs, including monospecific antibody-sialidase constructs^12–14^ and bispecific T cell engager-sialidase fusions^15^. Across preclinical models, these approaches enhanced antitumor activity through effects spanning tumor, myeloid, and lymphoid compartments^10–12,14–16^.

Whether sialylation on human T cells constitutes a checkpoint comparable to tumor-cell hypersialylation is far less clear. In murine T cells, activation, differentiation, and aging are accompanied by broad changes in surface sialylation^17,18,19^, and enzymatic desialylation of T cells or antigen-presenting cells can enhance antigen-driven responses^20–23^. These findings establish the functional relevance of murine T cell sialylation, however comparative work suggests that the regulation and functional consequences of T cell sialylation may differ between mice and humans^18^. It therefore remains unclear whether surface sialylation directly limits human T cell responses and whether this effect varies across subsets.

This question is particularly relevant to bispecific T cell engagers (TCEs), which redirect T cells to cancer cells by co-engaging a tumor antigen and CD3ε independently of peptide-major histocompatibility complex (pMHC) recognition^24^. TCEs are clinically established in several hematologic malignancies and are being actively developed for solid tumors, but their efficacy depends not only on target expression but also on the composition and functional state of the available T cell pool^25,26^. Both tumor-associated and peripheral T cells can contribute to TCE responses^27–30^, and clinical responses have been linked to contributions from naïve CD8^+^ T cells as sources of highly expandable effectors^31^. Yet TCE responses are often dominated by memory T cells, consistent with the greater costimulatory requirements and stricter activation thresholds of naïve cells^32,33^. A sialoglycan-dependent restraint could therefore be particularly relevant in naïve and other less-differentiated T cell populations. Preclinical studies have shown that sialidase administration increased TCE-mediated anti-tumor activity in preclinical tumor models^15^, but they did not resolve T cell dependent mechanisms.

To address this gap, we used *Arthrobacter ureafaciens* sialidase (AUS) to remove surface sialic acids from T cells isolated from healthy-donor peripheral blood mononuclear cells (PBMCs) and assessed TCE-induced activation, proliferation, cytokine production, and tumor-cell killing *in vitro*. We then extended the analysis to primary chronic lymphocytic leukemia (CLL) PBMC cultures treated with AUS and the CD20-directed TCE glofitamab. CLL provided a clinically relevant autologous setting in which malignant B cells coexist with dysfunctional T cells and responses to CD20-directed TCEs remain suboptimal^34–37^. In the CLL samples, single-cell RNA sequencing (scRNA-seq) combined with lectin-based cellular indexing of transcriptomes and epitopes by sequencing (CITE-seq) linked T cell subset identity and treatment-induced transcriptional states to lectin-defined cell-surface glycan signatures. Across both systems, naïve CD4^+^ and CD8^+^ T cells and, to a lesser extent, central memory T cells showed elevated α2,3-sialylation signatures and the clearest responses to desialylation. In healthy-donor cultures, desialylation enhanced activation, proliferation, cytokine production, and TCE-mediated tumor-cell killing; in CLL samples, it enhanced cytotoxic effector transcriptional programming. CD43 emerged as a major carrier of α2,3-linked sialoglycans and a partial contributor to sialidase responsiveness. Together, these findings identify T cell surface sialylation as a subset-specific restraint on TCE responses and provide a rationale for testing sialidase-TCE combinations designed to increase the contribution of less-differentiated T cells.

## Results

### Sialidase treatment enhances activation of human T cells

On mammalian glycoproteins, terminal sialic acids are commonly attached to N- and O-linked glycans through α2,3 or α2,6 linkages^3^, which can be profiled by lectin staining. To assess linkage-defined sialylation across human immune cells, freshly isolated PBMCs from healthy donors (HDs) were stained with the lectins *Maackia amurensis* lectin II (MAL II), which recognizes α2,3 sialic acids on O-glycans, and *Sambucus nigra* agglutinin (SNA), which preferentially binds to sialic acids on terminal galactoses in an α2,6 linkage^38^. CD4□ and CD8□ T cells displayed substantially higher MAL II signal intensity than other PBMC subsets, indicating particularly prominent α2,3-sialylation in peripheral-blood T cells (**Fig. 1A, Fig. S1A**). In contrast, monocytes showed the highest SNA signal, consistent with higher α2,6-sialylation (**Fig. 1A**).

**Fig. 1:**
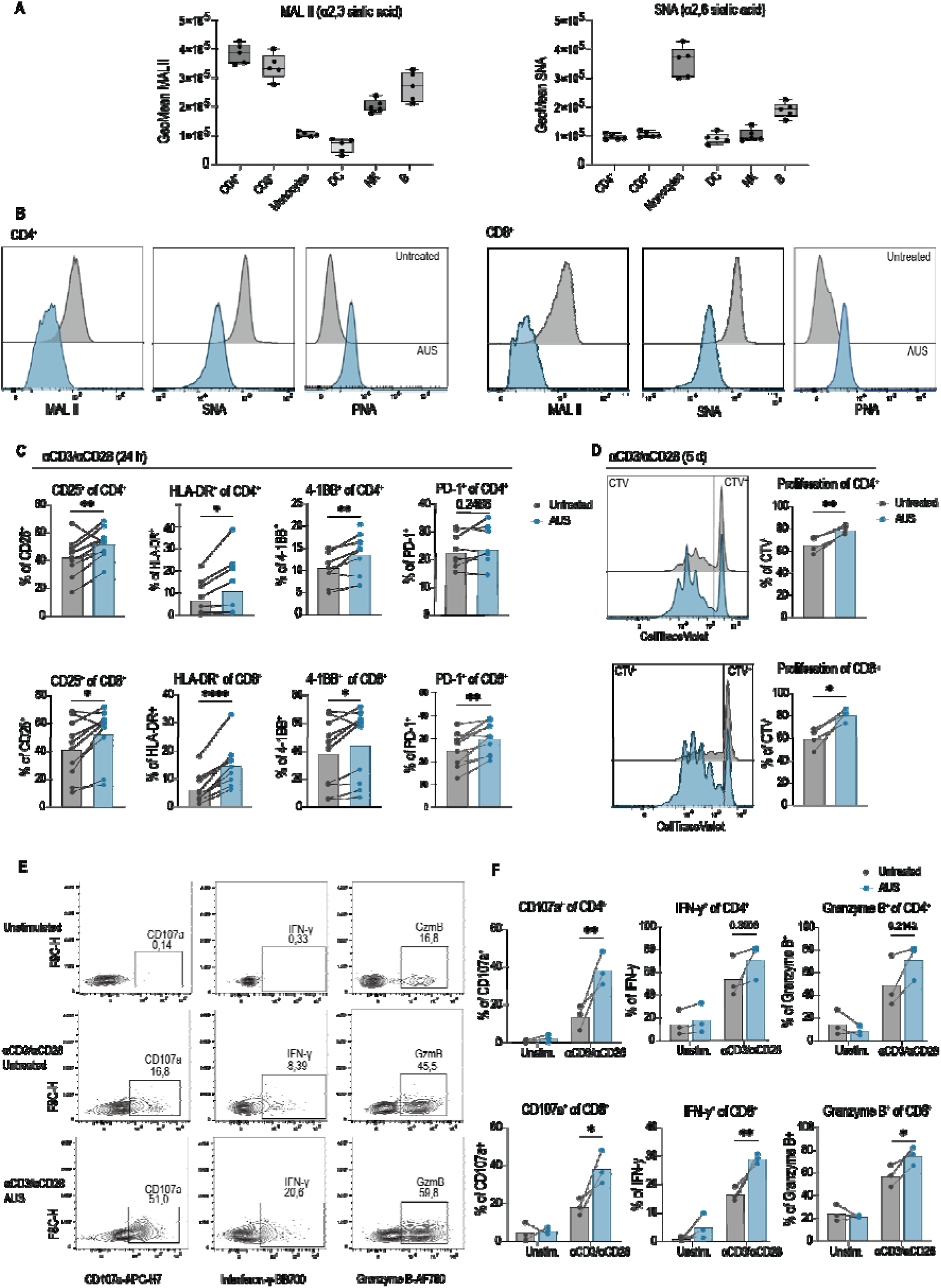
Sialidase treatment enhances human T cell activation and effector function. **A)** Flow-cytometric lectin staining of HD PBMCs with MAL II and SNA. GeoMean fluorescence intensity is shown for the indicated immune-cell subsets (*n* = 5 donors). B, B cells; DC, dendritic cells; NK, natural killer cells. **B)** Representative histograms of MAL II, SNA, and PNA lectin binding on HD CD4^+^ and CD8^+^ T cells after AUS treatment. **C)** Expression of activation markers measured by flow cytometry after stimulation of CD4^+^ and CD8^+^ T cells with αCD3/αCD28 for 24 hours ± 10 U/mL AUS. Pooled data with *n* = 8 donors from *n* = 3 experiments are shown. **D)** Representative histograms and frequencies of proliferated CTV^−^ CD4^+^ and CD8^+^ T cells stimulated with αCD3/αCD28 for 5 days ± 10 U/mL AUS (*n* = 4 donors, *n* = 2 experiments). **E)** Representative contour plots showing CD107a, GzmB, and IFN-γ staining in CD8^+^ T cells stimulated with αCD3/αCD28 ± 10 U/mL AUS for 3 days. **F)** Quantification of CD107a^+^, GzmB^+^, and IFN-γ in CD4 and CD8 T cells stimulated with αCD3/αCD28 ± 10 U/mL AUS for 3 days (*n* = 3 donors). Statistical significance was assessed by two-tailed paired Student’s t test for **C** and **D** and by two-way ANOVA with Holm–Sidak’s multiple-comparison test for **F**. \**p* < 0.05, \*\**p* < 0.01, \*\*\**p* < 0.001, \*\*\*\**p* < 0.0001. Abbreviations: AUS, *Arthrobacter ureafaciens* sialidase; CTV, CellTrace Violet; GeoMean, geometric mean; GzmB, granzyme B; HD PBMCs, healthy-donor peripheral blood mononuclear cells; IFN-γ, interferon-γ; MAL II, *Maackia amurensis* lectin II; PNA, peanut agglutinin; SNA, *Sambucus nigra* agglutinin.

We next asked whether enzymatic desialylation alters T cell function. Treatment with *Arthrobacter ureafaciens* sialidase (AUS), a broad-spectrum sialidase, markedly reduced MAL II and SNA binding on HD PBMC-derived T cells while increasing peanut agglutinin (PNA) staining, confirming efficient removal of α2,3- and α2,6-linked terminal sialic acids and exposure of underlying core-1 O-glycan structures (**Fig. 1B**, **Fig. S1B**).

Upon T cell receptor (TCR) and CD28 stimulation with agonistic anti-CD3 and anti-CD28 antibodies (αCD3/αCD28), desialylation increased expression of the activation markers CD25, HLA-DR, and 4-1BB in both CD4^+^ and CD8^+^ T cells after 24 h of culture, with increased PD-1 expression in CD8^+^ T cells (**Fig. 1C**). Desialylation further enhanced T cell proliferation after 5 days of culture, as shown by CellTrace Violet (CTV) dilution (**Fig. 1D**). In addition, sialidase-treated CD4^+^ and CD8^+^ T cells showed increased degranulation (CD107a), and CD8^+^ T cells produced higher levels of effector molecules, including interferon-γ (IFN-γ) and granzyme B (**Fig. 1E**,**F**)^39^. Notably, these effects were observed only in stimulated T cells, indicating that sialidase alone does not activate resting T cells (**Fig. 1E**,**F**).

Because sialylated glycans are described to attenuate CD28 costimulation by acting as alternative CD28 ligands^23^, we tested whether AUS-enhanced activation was primarily attributable to relief from this CD28-dependent inhibitory mechanism. AUS-enhanced activation was retained when T cells were stimulated with plate-bound αCD3 in the absence of CD28 costimulation, indicating that CD28 engagement was not required for the effect (**Fig. S2A**). AUS also enhanced activation after phorbol 12-myristate 13-acetate (PMA)/ionomycin stimulation, which bypasses TCR/CD3 ligation and directly activates downstream PKC- and calcium-dependent pathways (**Fig. S2B**). Thus, sialidase-mediated enhancement is not solely explained by release from CD28-dependent inhibition or by effects limited to receptor-proximal TCR/CD3 engagement.

Together, these findings indicate that surface sialylation restrains human T cell activation across distinct stimulation contexts, prompting us to test whether desialylation could also enhance TCE-mediated activation and tumor-cell killing.

### T cell desialylation enhances T cell engager-mediated activation and tumor-cell killing

To assess whether desialylation enhances T cell function in a therapeutically relevant TCE setting, HD PBMC-derived T cells were co-cultured with CD20^+^ Raji B cell lymphoma target cells in the presence or absence of AUS. Co-cultures were stimulated with glofitamab, a CD20-directed bispecific TCE approved for B cell malignancies^40^ (**Fig. 2A**). After overnight co-culture, CD4^+^ and CD8^+^ T cells exposed to AUS showed increased expression of CD25 and CD69 compared with untreated control T cells (**Fig. 2B**).

**Fig. 2:**
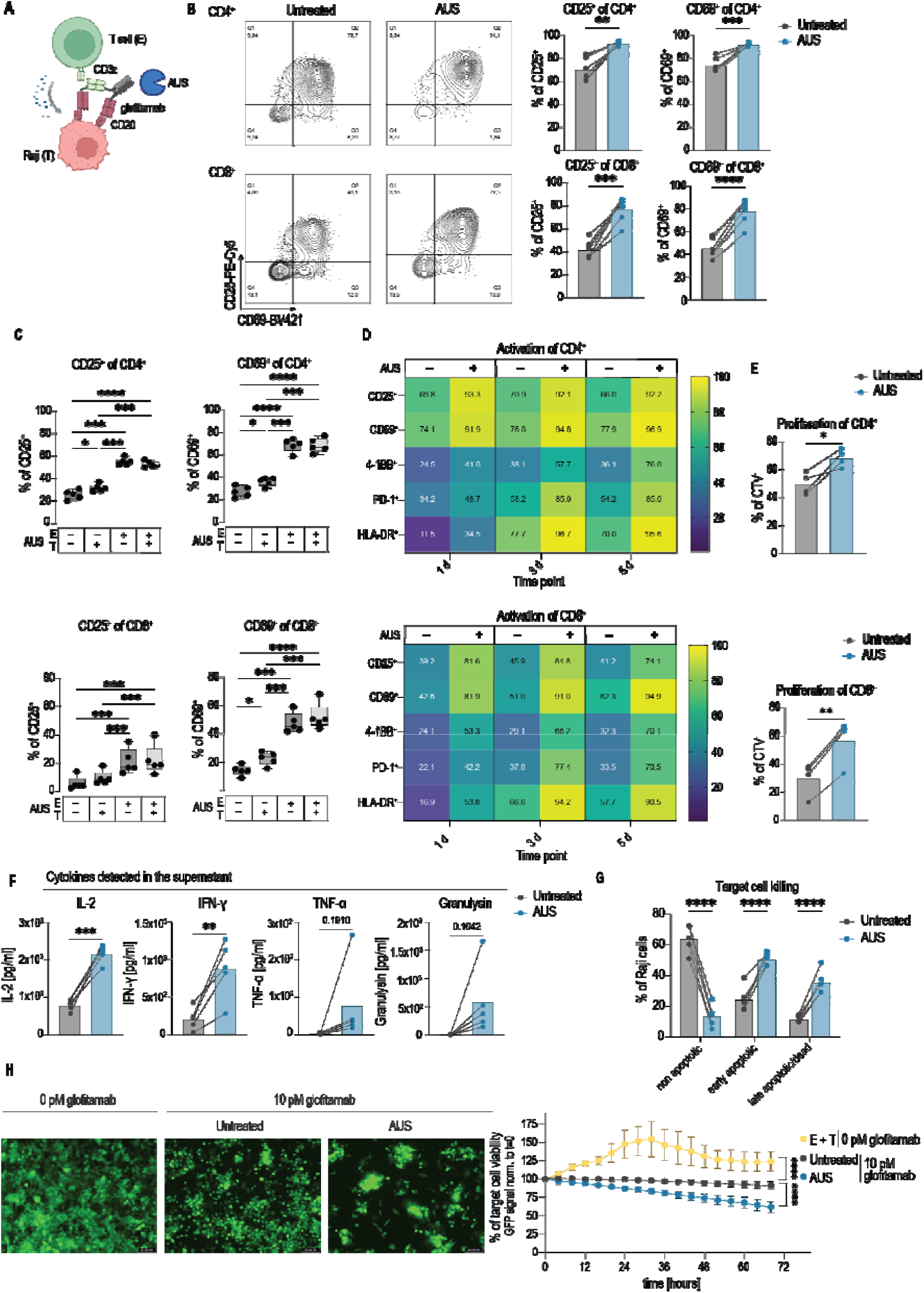
T cell desialylation enhances glofitamab-mediated activation and tumor cell killing. **A)** Schematic of the Raji/glofitamab co-culture T cell activation assay using AUS-treatment. **B)** Representative contour plots and frequencies of CD25^+^ and CD69^+^ CD4^+^ and CD8^+^ T cells after 24 h co-culture with Raji cells and 0.1 pM glofitamab ± 10 U/mL AUS; *n* = 6 donors, **C)** Frequencies of CD25^+^ and CD69^+^ CD4^+^ and CD8^+^ T cells after 24 h co-culture with Raji target cells and 0.1 pM glofitamab following compartment-specific desialylation. Effector T cells (E) and/or Raji target cells (T) were pre-treated separately with AUS, washed to remove residual enzyme, and then combined for co-culture. “+” indicates AUS treatment; “-” indicates untreated control; *n* = 5 donors. **D)** Time course of activation-marker expression in CD4^+^ and CD8^+^ T cells on days 1, 3, and 5 after stimulation with 0.1 pM glofitamab ± 10 U/mL AUS. Frequencies of CD25^+^, CD69^+^, 4-1BB^+^, PD-1^+^, and HLA-DR^+^ T cells are shown; *n* = 6 donors. **E)** Frequencies of proliferated CTV^−^ CD4^+^ and CD8^+^ T cells after stimulation with glofitamab for 5 days ± 10 U/mL AUS; *n* = 4 donors from 2 independent experiments. **F)** Cytokines and effector molecules measured in co-culture supernatants after 24 hours of Raji/T cell/glofitamab stimulation ± 10 U/mL AUS; *n* = 5 donors. **G)** Flow cytometric assessment of caspase-3 activity in Raji-dTomato-Luc2 target cells after 48-hour co-culture with T cells ± 10 U/mL AUS and 10 nM glofitamab; *n* = 5 donors. Non-apoptotic cells were defined as Zombie NIR^−^/Caspase-3^−^, early apoptotic as Zombie NIR^−^/Caspase-3^+^ and late apoptotic as NIR^+^/Caspase-3^−/+^. **H)** Left: Representative GFP images of A375-GFP-CD20 melanoma target cells after 68 h of co-culture with T cells and 10 pM glofitamab ± 10 U/mL AUS. Right: Quantification of GFPO target cells by real-time IncuCyte imaging. Data are shown as meanO±OSD; *n* = 3 donors with 3 technical replicates each. Curves show nonlinear regression fits for samples treated with 10 pM glofitamab. Statistical significance was assessed by two-tailed paired Student’s t test for **B** and **F**, two-way ANOVA with Tukey correction for **C**, two-way ANOVA with Holm–Sidak correction for **E**, and one-way ANOVA with Holm–Sidak correction for **G** and **H**. For **H**, statistical significance was determined at 68 hours. \**p* < 0.05, \*\**p* < 0.01, \*\*\**p* < 0.001, \*\*\*\**p* < 0.0001. Abbreviations: AUS, *Arthrobacter ureafaciens* sialidase; CTV, CellTrace Violet; GFP, green fluorescent protein; HD, healthy donor; TCE, T cell engager.

Because tumor-associated sialylation can impair T cell cytotoxicity^41,22^, we asked whether this enhancement was driven by desialylation of T cells, target cells, or both. Separate AUS pre-treatment of the effector and target compartments showed that target-cell desialylation produced a modest increase in T cell activation, whereas T cell-directed desialylation had the strongest effect, significantly upregulating CD25 and CD69 expression (**Fig. 2C**). Simultaneous desialylation of T cells and target cells did not further increase activation beyond T cell-directed desialylation alone (**Fig. 2C**).

We then applied sialidase directly to the co-culture to approximate a therapeutic setting, in which effector and target compartments would be exposed to sialidase simultaneously. Longitudinal assessment over a 5-day Raji/T cell co-culture showed that sialidase enhanced both early and late glofitamab-mediated activation of CD4^+^ and CD8^+^ T cells, with upregulation of CD25, CD69, 4-1BB, PD-1, and HLA-DR (**Fig. 2D**). Desialylation also increased proliferation of glofitamab-treated CD4^+^ and CD8^+^ T cells (**Fig. 2E**) and enhanced release of interleukin-2 (IL-2) and IFN-γ, with non-significant trends for tumor necrosis factor-α (TNF-α) and granulysin (**Fig. 2F**).

This enhanced activation was accompanied by increased target-cell killing. Flow-cytometric analysis of dTomato-expressing Raji target cells revealed a higher proportion of apoptotic and dead cells in AUS-treated co-cultures than in untreated controls (**Fig. 2G**, **Fig. S3A–D**), whereas sialidase alone did not induce target-cell death (**Fig. S3E**). To corroborate these findings in a distinct target-cell model and enable longitudinal monitoring of tumor-cell killing, we used A375 melanoma cells engineered to express green fluorescent protein (GFP) and CD20. In this live-cell imaging assay, sialidase treatment accelerated loss of GFP signal over time, further supporting that AUS enhances glofitamab-mediated tumor-cell killing (**Fig. 2H**).

Control experiments showed that enhanced activation required enzymatically active sialidase and was recapitulated by selective α2,3-desialylation. Specifically, heat-inactivated AUS neither reduced MAL II/SNA binding nor enhanced glofitamab-induced CD25 or CD69 expression (**Fig. S4A**,**B**). Conversely, *Streptococcus pneumoniae* α2,3-sialidase (SPS) reduced MAL II but not SNA binding and enhanced CD25 and CD69 expression to levels comparable to AUS (**Fig. S4C**,**D**)^42^, indicating that selective desialylation of α2,3-linked sialoglycans is sufficient to enhance TCE-induced activation under these conditions.

We next tested whether two known immune-regulatory pathways could account for the early activation-enhancing effect of AUS. Sialic acid-binding immunoglobulin-like lectins (Siglecs) can mediate sialic acid-dependent immune suppression. Siglec-7 and Siglec-9 have been implicated in suppression of human intratumoral T cells^43,44^, raising the possibility that AUS enhanced activation by disrupting this inhibitory axis. However, in HD-PBMCs only a small fraction of T cells express Siglecs at low levels^43,44^. Consistently, Siglec-7 and Siglec-9 expression on HD PBMC-derived T cells remained largely undetectable during the 24-hour activation window used for CD25 and CD69 analysis, with only low-level Siglec-9 expression emerging after 3 and 5 days of culture (**Fig. S5A**,**B**). Consistent with this expression pattern, antibody blockade of Siglec-7 and Siglec-9 did not alter CD25 or CD69 expression compared with the isotype control (**Fig. S5C**), arguing against Siglec-7/9 engagement as a primary explanation for sialidase-enhanced activation. We next tested whether costimulation through the CD80/CD86-CD28 axis contributed to the sialidase effect, because Raji cells express both CD80 and CD86 and can engage CD28 during glofitamab-mediated T cell activation^23,45^. Sialidase-enhanced activation was retained with CRISPR/Cas9-generated Raji CD80/CD86 double-knockout cells (**Fig. S5D–F**), confirming that CD28-mediated costimulation was not required in this model.

Having established a predominantly T cell-directed effect in the Raji/glofitamab system, we next asked whether T cell desialylation also enhanced activation in additional TCE formats. In co-cultures using CD19□ Raji cells with blinatumomab, DLL3□ SHP-77 cells with tarlatamab, and EpCAM□ OVCAR-8 cells with a CD3×EpCAM TCE, increased CD25 and CD69 expression in CD4^+^ and CD8^+^ T cells was consistently driven by T cell-directed desialylation (**Fig. S6A–C**). By contrast, modulation of tumor cell sialylation produced less pronounced and model-dependent effects. Lectin profiling of the corresponding target-cell lines revealed differences in both baseline sialylation and the extent of AUS-mediated desialylation, which may help explain this variability (**Fig. S6D**).

Together, these findings indicate that sialoglycans on T cells restrain TCE-induced activation across distinct engager formats, target antigens, and tumor-cell contexts. We therefore next asked whether sialidase treatment could also enhance glofitamab responses in primary samples from patients with CLL, a disease in which malignant CD20^+^ B cells coexist with dysfunctional autologous T cells and responses to CD20-directed TCEs remain suboptimal^34–37^.

### Sialidase enhances glofitamab-induced effector programming in CLL T cells

To determine whether desialylation alters glofitamab-induced T cell programs in primary CLL samples, we profiled *ex vivo* CLL PBMC cultures by scRNA-seq coupled with lectin-based CITE-seq, linking treatment-induced transcriptional states to lectin-defined cell-surface glycan signatures within the same single-cell dataset (**Fig. 3A**).

**Fig. 3:**
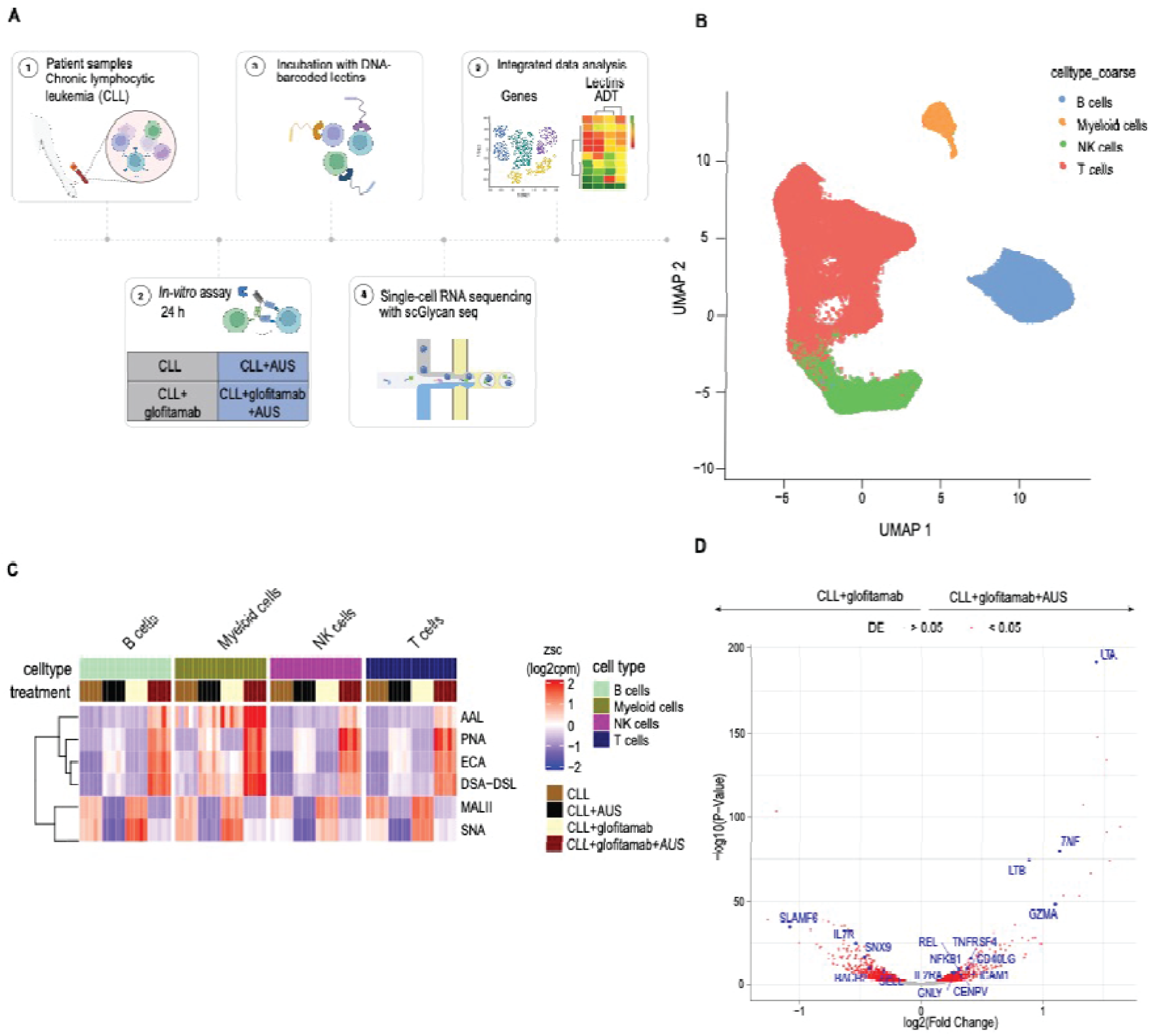
Sialidase enhances glofitamab-induced effector programming in CLL T cells. **A)** Schematic of the *ex vivo* CLL single-cell glycan-profiling workflow. PBMCs from patients with CLL were cultured overnight with or without 1 pM glofitamab and in the presence or absence of 10 U/mL AUS, followed by CD19-based enrichment, antibody and lectin staining, flow-cytometric sorting, and downstream scRNA-seq coupled to lectin-based CITE-seq. Gene-expression profiles were integrated with lectin-derived ADT signals. **B)** UMAP of all cells retained after quality control, based on gene-expression profiles and annotated into B cells, T cells, NK cells, and myeloid cells using the Human Cell Atlas^46^. All experimental conditions are included; each dot represents one cell. **C)** Heatmap of normalized lectin-derived ADT signals across major immune-cell populations and treatment conditions. Values are shown as row-wise z scores of normalized ADT abundance, capped at −2 and 2 for visualization. **D)** Volcano plot showing differentially expressed genes in T cells comparing combined treatment with CLL + glofitamab + AUS to CLL + glofitamab alone. Positive log2 fold changes indicate higher expression after combined glofitamab and AUS treatment; negative values indicate higher expression after glofitamab alone. Genes passing absolute log2 fold change > 0.3 and Benjamini–Hochberg FDR-adjusted P < 0.05 are highlighted. Abbreviations: AAL, *Aleuria aurantia* lectin; ADT, antibody/affinity-derived tag; AUS, *Arthrobacter ureafaciens* sialidase; CITE-seq, cellular indexing of transcriptomes and epitopes by sequencing; CLL, chronic lymphocytic leukemia; DSA, *Datura stramonium* agglutinin; ECA, *Erythrina cristagalli* agglutinin; FDR, false discovery rate; MAL II, *Maackia amurensis* lectin II; NK, natural killer; PBMCs, peripheral blood mononuclear cells; PNA, peanut agglutinin; scRNA-seq, single-cell RNA sequencing; SNA, *Sambucus nigra* agglutinin; UMAP, uniform manifold approximation and projection.

PBMCs from patients with CLL (*n* = 5, **Supplementary Table 1**) were cultured overnight with or without glofitamab in the presence or absence of AUS (**Fig. 3A**). HI-AUS served as the control in samples not treated with AUS.

Because CLL PBMCs were dominated by malignant CD19^+^ B cells, which showed variable CD20 expression across donors, CD19^−^ immune cells were enriched after culture and recombined with a defined CD19^+^ fraction before flow-cytometric sorting for single-cell profiling (**Fig. S7A–D**). After quality control, 80529 cells were retained for analysis. Gene expression-based clustering and annotation against the Human Cell Atlas^46^ identified T cells, B cells, natural killer (NK) cells, and a minor myeloid population (**Fig. 3B**).

Lectin-CITE-seq resolved treatment-induced changes in surface glycan motifs across immune-cell populations. In samples not treated with AUS, major immune populations, including T cells, showed robust binding of MAL II and SNA (**Fig. 3C**). AUS treatment reduced MAL II and SNA signals and increased binding of PNA, *Erythrina cristagalli* agglutinin (ECA), and *Datura stramonium* agglutinin (DSA), consistent with desialylation and increased accessibility of underlying glycan structures. By contrast, *Aleuria aurantia* lectin (AAL) binding was most prominent after combined glofitamab and AUS treatment, suggesting additional activation-associated changes in AAL-reactive glycan structures (**Fig. 3C**). These findings corroborate the flow-cytometric lectin staining data from HD PBMC-derived T cells (**Fig. 1C**) and support the use of lectin-CITE-seq to monitor desialylation at single-cell resolution.

We next asked how desialylation altered the transcriptional response of disease-associated T cells to glofitamab. Glofitamab alone induced an activation and effector signature^47,48^ characterized by upregulation of activation-associated genes (*IL2RA, TNFRSF9, CTLA4*), cytotoxic effector genes (*PRF1, GZMB, LTA*), and proinflammatory cytokine genes (*TNF, IFNG*) (**Fig. S8**). Compared with glofitamab alone, combined treatment with glofitamab and AUS further increased expression of effector- and activation-associated genes, including *LTA*, *TNF, LTB, TNFRSF4, CD40LG,* and *GNLY* (**Fig. 3D**). By contrast, exhaustion-associated genes (*PDCD1, HAVCR2, LAG3*) were not detectably increased in this comparison, indicating that short-term desialylation enhanced effector programming without acutely amplifying exhaustion-associated transcripts (**Fig. S8**).

Together, these data show that sialidase treatment alters lectin-defined surface glycan signatures in primary CLL samples and strengthens the glofitamab-induced cytotoxic effector transcriptional program in autologous T cells. This prompted us to ask whether the transcriptional response to desialylation was broadly distributed across T cells or concentrated within specific subsets.

### Sialidase preferentially reprograms highly sialylated less-differentiated CLL T cells

To define which T cell subsets were most responsive to sialidase treatment, we re-clustered T cells from the CLL single-cell dataset and resolved 12 transcriptionally distinct subpopulations using published signatures and canonical marker genes^46,49^ (**Fig. 4A**, **Fig. S9A**). Glofitamab-containing conditions gave rise to activated counterparts of different CD4^+^ and CD8^+^ T cell subsets, which formed distinct UMAP clusters (**Fig. 4B**).

**Fig. 4:**
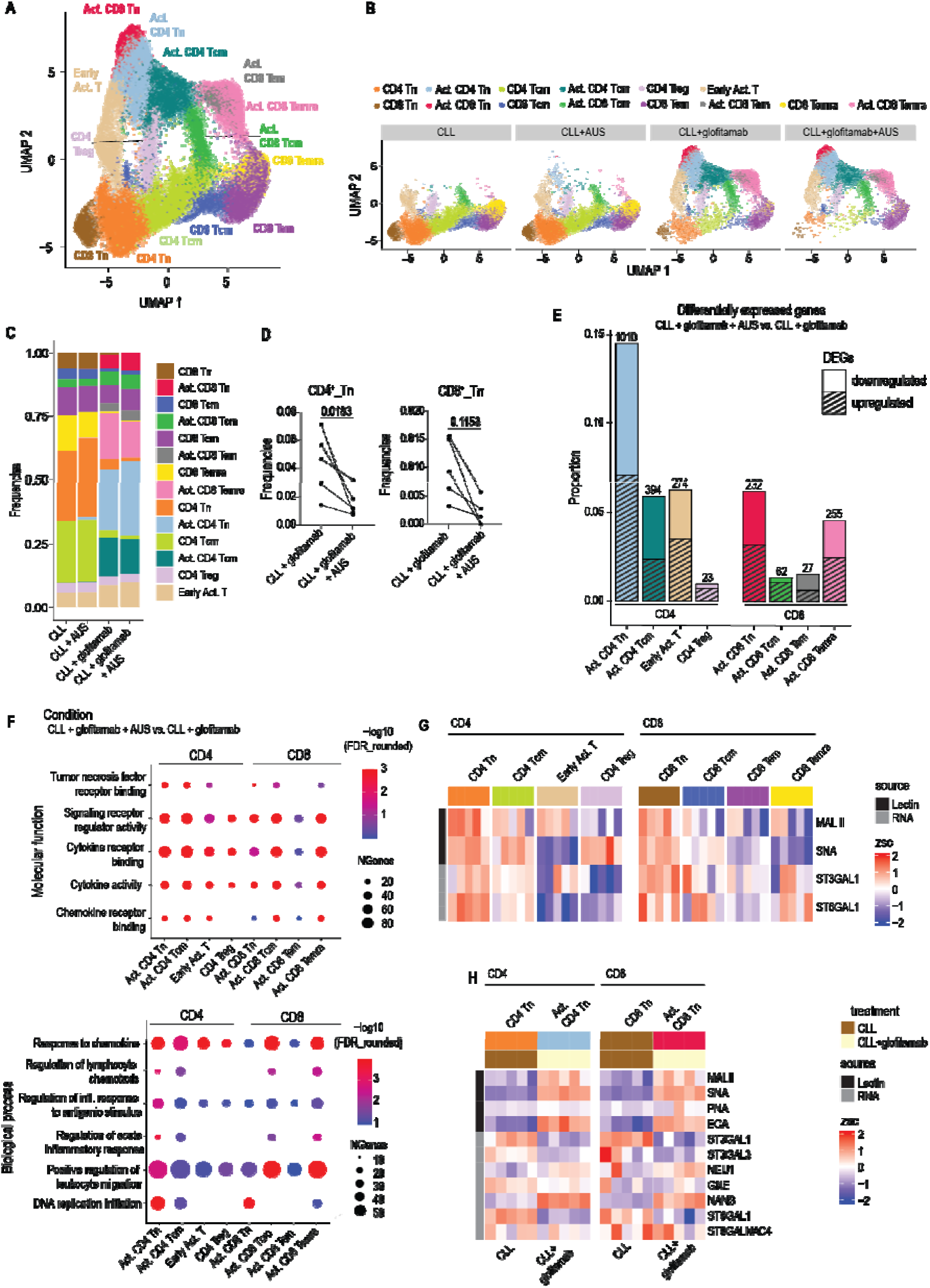
Sialidase preferentially reprograms highly sialylated less-differentiated CLL T cells. **A)** UMAP of re-clustered T cells from the CLL single-cell dataset, annotated into transcriptionally distinct T cell subsets using canonical marker genes and published T cell signatures^46,49^. **B)** UMAPs of annotated T cell subsets faceted by treatment conditions: CLL, CLL + AUS, CLL + glofitamab, CLL + glofitamab + AUS. **C)** Frequencies of annotated T cell subsets across the 4 treatment conditions. **D)** Frequencies of naïve CD4^+^ and CD8^+^ T cells in CLL + glofitamab versus CLL + glofitamab + AUS. Paired samples from the same donor are connected by lines and statistics are reported as FDR adjusted p values. **E)** Proportions of DEGs within each T cell subset comparing CLL + glofitamab + AUS with CLL + glofitamab. Bar height indicates the proportion of expressed genes that were differentially expressed within each subset; numbers above bars indicate the absolute number of DEGs. Hatched and non-hatched segments indicate upregulated and downregulated genes, respectively. **F)** GO enrichment analysis of genes upregulated in CLL + glofitamab + AUS compared with CLL + glofitamab. Selected molecular-function and biological-process GO terms are shown for CD4^+^ and CD8^+^ T cell subsets. Dot size indicates the number of DEGs contributing to each GO term, and color indicates −log10(FDR), FDR values < 0.001 were set to 0.001 for visualization (FDR rounded). **G)** Heatmap of lectin-derived ADT signals for MAL II and SNA together with RNA expression of *ST3GAL1* and *ST6GAL1* across selected CD4O and CD8O T cell subsets in the untreated condition. **H)** Heatmap of lectin-derived ADT signals and RNA expression of genes involved in sialoglycan biosynthesis in naïve CD4O and CD8O T cells from untreated CLL cultures and their activated counterparts after glofitamab stimulation. For **G** and **H**, RNA expression values are shown as row-wise z scores of normalized log2(cpm). For lectin-derived ADT signals, normalized ADT abundance was averaged across cells within each subset, treatment condition, and donor before row-wise z scoring. Z scores were capped at −2 and 2 for visualization. Abbreviations: ADT, antibody/affinity-derived tag; AUS, *Arthrobacter ureafaciens* sialidase; CLL, chronic lymphocytic leukemia; DEG, differentially expressed gene; FDR, false discovery rate; GO, Gene Ontology; MAL II, *Maackia amurensis* lectin II; Tcm, central memory T cells; Tem, effector memory T cells, Temra, terminally differentiated effector memory T cells re-expressing CD45RA; Tn, naïve T cells, Treg, regulatory T cells, SNA, *Sambucus nigra* agglutinin; UMAP, uniform manifold approximation and projection.

Quantification of subset frequencies across treatments showed that combined glofitamab and AUS reduced the frequency of resting CD4^+^ naïve T cells, with a similar non-significant trend in CD8^+^ naïve T cells, compared with glofitamab alone (**Fig. 4C,D**, **Fig. S9B**). These changes were consistent with activation-linked redistribution of the naïve T cell compartment.

Differential gene expression analysis comparing combined glofitamab and AUS treatment with glofitamab alone showed that the transcriptional response to sialidase treatment was strongly subset-dependent. When normalized to the number of genes detected within each subset, activated naïve CD4^+^ and activated naïve CD8^+^ T cells displayed the highest differentially expressed gene (DEG) proportions within their respective lineages, with central memory subsets also exhibiting prominent remodeling (**Fig. 4E**, **Fig. S10**). Activated CD8^+^ Temra cells showed a sizable absolute number of DEGs, although their normalized DEG proportion was lower than that of activated naïve CD8^+^ T cells. Thus, the strongest glycan-linked transcriptional response was observed in less-differentiated T cell states, while activated CD8^+^ Temra cells represented an additional responsive population.

Gene Ontology enrichment analysis of genes upregulated by combined glofitamab and AUS treatment relative to glofitamab alone showed that sialidase strengthened cytokine-, chemokine-, and tumor necrosis factor receptor-related programs most prominently in activated naïve and central memory CD4^+^ T cells and activated naïve CD8^+^ T cells (**Fig. 4F**). These responsive subsets also exhibited enrichment of gene sets associated with cytokine response, DNA replication initiation, and leukocyte migration, consistent with enhanced activation, proliferative signaling, and migration-associated transcript remodeling. In line with these enriched programs, activated naïve CD4^+^ and CD8^+^ T cell states downregulated quiescence-associated transcripts, including *SELL*, *IL7R*, and *BACH2*, while upregulating effector- and activation-associated transcripts, such as *LTA*, *LTB*, *IFNG*, *TNF*, *TNFRSF4*, and *TNFRSF18* (**Fig. S10A**,**B**). *CXCR3* expression was increased most clearly in activated naïve CD4^+^ T cells, with a weaker trend in activated naïve CD8^+^ T cells, consistent with migration-associated transcriptional remodeling^50,51^ (**Fig. S10A**,**B**). In contrast, Tregs and most effector-memory subsets showed limited remodeling, although CD8^+^ Temra cells retained detectable enrichment of selected inflammatory and effector-associated programs (**Fig. 4F**).

Collectively, combined glofitamab and AUS treatment elicited heterogeneous, subset-specific transcriptional responses in CLL T cells. These findings prompted us to examine whether subset-specific responsiveness to desialylation corresponded to differences in cell-surface sialylation.

Having established that lectin-CITE-seq resolved treatment-associated changes in surface glycan signatures across immune lineages, we next leveraged this approach to compare baseline sialylation across defined T cell subsets from CLL samples. Guided by prior reports of elevated sialylation in murine naïve T lymphocytes ^17–19,52^, we asked whether naïve human T cells from CLL patients similarly displayed elevated lectin-defined sialylation at baseline. In untreated CLL PBMC cultures, naïve CD4^+^ and CD8^+^ T cells showed stronger MAL II and SNA binding than memory subsets, regulatory T (Tregs) cells, and Temra cells, indicating higher α2,3- and α2,6-sialylation in the naïve compartment (**Fig. 4G**). In line with surface lectin staining, naïve T cells also expressed higher levels of the α2,3-sialyltransferase *ST3GAL1* and the α2,6-sialyltransferase *ST6GAL1*^53^ (**Fig. 4G**, **Fig. S11A**,**B**). Together, these findings indicate that less-differentiated T cells in CLL samples are characterized by stronger lectin-defined sialylation signatures along with increased expression of sialylation-associated enzymes than more differentiated subsets.

We next asked how this profile differed between naïve and activated T cell states after glofitamab stimulation, using a curated gene signature of sialylation-associated enzymes^16^. Glofitamab-mediated activation of naïve CD4^+^ and CD8^+^ T cells was associated with downregulation of α2,3- and α2,6-sialyltransferases, including *ST3GAL1*, *ST3GAL3*, *ST6GAL1*, and *ST6GALNAC4,* together with increased expression of the sialic acid-removing neuraminidase *NEU1* (**Fig. 4H**, **Fig. S11C**,**D**). Together, these changes suggest an activation-associated shift in the sialylation machinery of naïve CLL T cells, consistent with previous work on T cell sialylation during activation and differentiation^18,53,54^.

Although baseline MAL II and SNA binding broadly corresponded to sialyltransferase expression (**Fig. 4G**), activated T cells showed higher signals for multiple lectins, including MAL II, SNA, PNA, and ECA. Because activation induces blastogenesis, these increases may partly reflect greater cell size and therefore cannot be interpreted as evidence of increased glycan density^55^.

Taken together, these findings reveal marked heterogeneity in lectin-defined sialylation profiles and transcriptional responsiveness to AUS across T cell subsets in CLL samples. Naïve and central memory T cells showed the clearest correspondence between elevated baseline lectin signals and responsiveness to combined glofitamab and AUS treatment, whereas activated CD8^+^ Temra cells represented a notable exception.

### Naïve T cell α2,3-sialoglycan remodeling implicates CD43 in sialidase responsiveness

Given the preferential responsiveness of naïve T cells in the CLL samples, we next tested whether this subset bias was preserved in the healthy-donor T cell-Raji co-culture model using flow cytometry. Consistent with the patient-derived data, AUS-mediated enhancement of glofitamab-induced activation was strongly subset-dependent: naïve T cells (Tn, CD45RA□CCR7□) showed the most pronounced increase in CD25 and CD69 expression, followed by central memory T cells (Tcm, CD45RA□CCR7□), whereas effector memory T cells (Tem, CD45RA□CCR7□) and Temra cells (CD45RA□CCR7□) showed smaller responses (**Fig. 5A**,**B**). This responsiveness gradient aligned with baseline glycan features: naïve and central memory T cells displayed the strongest MAL II binding, consistent with higher α2,3-sialylation, a pattern similar to that observed in CLL samples, whereas SNA staining did not resolve comparable subset differences in α2,6-linked sialylation (**Fig. 5C**).

**Fig. 5:**
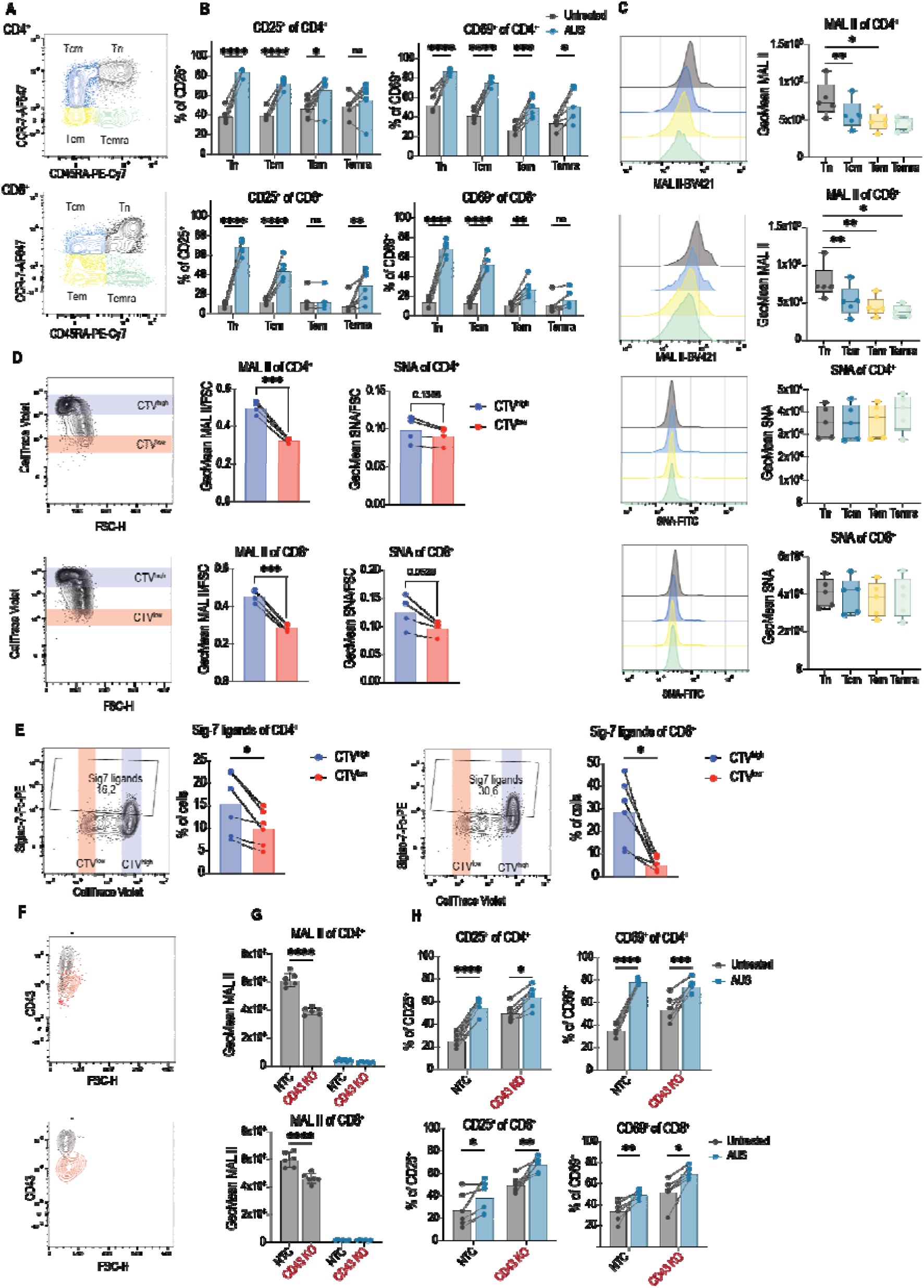
Naïve T cell α2,3-sialoglycan remodeling implicates CD43 in sialidase responsiveness. **A)** Representative flow-cytometric contour plots of CD4^+^ and CD8^+^ T cell differentiation subsets from HD PBMC-derived T cells based on CD45RA and CCR7 expression. Tn were defined as CD45RA^+^CCR7^+^, Tcm as CD45RA^−^CCR7^+^, Tem as CD45RA^−^CCR7^−^, and Temra cells as CD45RA^+^CCR7^−^. **B)** Frequencies of CD25^+^ and CD69^+^ cells among CD4^+^ and CD8^+^ T cell subsets after 24 h co-culture with Raji target cells and 0.1 pM glofitamab in the presence or absence of 10 U/ml AUS. “Untreated” denotes no AUS. Paired samples from the same donor are connected by lines. **C)** MAL II and SNA lectin binding on T cell subsets. Left, representative histograms; right, GeoMean fluorescence intensities of MAL II and SNA binding. Data shown as mean ± SD, *n* = 5 donors. **D)** MAL II and SNA lectin binding in naïve CD4O and CD8O T cells after 5-day stimulation with αCD3/αCD28. Lectin binding was normalized to cell size and is shown as GeoMean/FSC for early proliferative generations and late proliferative generations, defined by CTV dilution; *n* = 4 donors. **E)** Siglec-7-Fc binding in naïve CD4O and CD8O T cells activated with αCD3/αCD28 for 5 days. Left, representative contour plots; right, quantification of Siglec-7-Fc^+^ cells in undivided CTV^high^ and proliferated CTV^low^ populations; *n* = 5 donors. **F)** Representative contour plots showing CD43 expression in NTC and CD43 KO cells. **G)** MAL II binding in NTC and CD43 KO cells after co-culture with Raji target cells and 0.1 pM glofitamab ± 10 U/mL AUS. **H)** Frequencies of CD25^+^ and CD69^+^ cells among NTC and CD43 KO CD4^+^ and CD8^+^ T cells after Raji/glofitamab co-culture ± 10 U/mL AUS. Statistical significance was assessed by one-way ANOVA with Holm–Sidak correction for **C**, paired two-tailed Student’s t test for **D** and **E**, and two-way ANOVA with Holm–Sidak correction for **G** and **H**. Paired samples from the same donor are connected by lines where applicable. \**p*O<O0.05, \*\**p*O<O0.01, \*\*\**p*O<O0.001 and \*\*\*\**p*O<O0.0001. Abbreviations: AUS, *Arthrobacter ureafaciens* sialidase; CTV, CellTrace Violet; FSC, forward scatter; GeoMean, geometric mean fluorescence intensity; HD, healthy donor; KO, knockout; MAL II, *Maackia amurensis* lectin II; NTC, non-targeting control; PBMCs, peripheral blood mononuclear cells; SNA, *Sambucus nigra* agglutinin; Tcm, central memory T cells; Tem, effector memory T cells; Temra, terminally differentiated effector memory T cells re-expressing CD45RA; Tn, naïve T cells.

To characterize α2,3-sialoglycan remodeling during naïve T cell activation, we isolated naïve T cells, activated them for 5 days, and compared early and late proliferative generations defined by CTV dilution (**Fig. 5D**). Because activated T cells undergo blastogenesis, proliferating CTV^low^ cells showed increased forward scatter (FSC), indicating increased cell size (**Fig. S12A–D**). Raw lectin fluorescence can therefore reflect cell size and total per-cell signal as well as glycan density (**Fig. S12E**,**F**). To account for this, lectin signals were normalized to FSC (**Fig. S12G**,**H**). After normalization, MAL II binding showed a decrease in proliferating CTV^low^ cells, consistent with reduced α2,3-sialylation during activation, whereas SNA binding showed only a modest, nonsignificant reduction and PNA staining did not increase significantly (**Fig. 5D, Fig. S12G**,**H**). These findings suggest that human naïve T cell activation is associated most clearly with remodeling of α2,3-linked sialoglycans. In contrast, murine T lymphocytes have been reported to undergo predominantly α2,6-linked remodeling during effector memory differentiation^17,56,19^, indicating a divergence in glycan regulation between human and murine T cell activation, as described previously^18^.

We next asked which α2,3-sialylated O-glycoproteins might contribute to the sensitivity of less-differentiated T cells to AUS. CD43 emerged as a candidate because it is a major mucin-like T cell surface glycoprotein, a known ST3GAL-dependent substrate, and highly sialylated in naïve T cells^57^. It also regulates TCR-driven activation by modulating adhesion and receptor organization at the immunological synapse^57–59^, and its glycosylation shifts during T cell activation from highly sialylated core-1 O-glycans toward less-sialylated core-2 forms^57^.

To assess activation-associated remodeling of CD43-associated sialylated core-1 O-glycans, we used recombinant Siglec-7-Fc as a glycoform-sensitive probe. Unlike conventional CD43 staining, which cannot distinguish among differently glycosylated forms of CD43, Siglec-7-Fc detects the sialylated core-1 O-glycan form previously identified as a major Siglec-7 ligand^60^.

CTV-labeled T cells were activated with αCD3/αCD28, and Siglec-7-Fc binding was quantified across undivided CTV^high^ and proliferated CTV^low^ populations by flow cytometry, using AUS-treated cells as negative control to define the gate (**Fig. S13**). Siglec-7-Fc binding was high in undivided CTV^high^ cells and decreased significantly in proliferated CTV^low^ cells (**Fig. 5E**), consistent with activation-driven remodeling of sialic acid-dependent Siglec-7 ligands. Because Siglec-7-Fc is not CD43-specific, however, this readout supports but does not by itself prove CD43 glycan remodeling. We then tested whether CD43 contributed functionally to elevated α2,3-sialylation and to their responsiveness to AUS. To preserve a less-activated T cell state, CD43 was deleted in primary T cells by CRISPR/Cas9 before stimulation, with non-targeted sgRNAs serving as controls^61^. Sorted CD43 knockout and control cells were co-cultured with Raji target cells and stimulated with glofitamab in the presence or absence of AUS (**Fig. 5F**). CD43 knockout significantly reduced MAL II binding, supporting a role for CD43 as a major carrier of α2,3-linked sialoglycans in this setting (**Fig. 5G**). CD43-knockout cells also showed increased CD25 and CD69 expression after glofitamab stimulation alone compared with control cells, consistent with the reported lower activation threshold of CD43-deficient T cells^62^ (**Fig. 5H**). AUS-mediated enhancement of T cell activation was reduced but not abolished in CD43-knockout cells, indicating that CD43 contributes to sialidase-enhanced activation but is not the only relevant sialoglycoprotein.

Together, these findings suggest that activation of human naïve T cells is associated with reduced α2,3-sialylation and identify CD43 as a major carrier of α2,3-linked sialoglycans that contributes to AUS responsiveness. The persistence of AUS-enhanced activation after CD43 knockout indicates that additional sialoglycoproteins beyond CD43 contribute to the restraint of TCE-mediated T cell activation.

## Discussion

Our data identify T cell surface sialylation, particularly α2,3-linked sialoglycans, as a subset-specific restraint on human T cell activation and TCE responsiveness. Across HD T cell co-cultures and primary CLL samples, naïve T cells and, to a lesser extent, central memory T cells showed elevated α2,3-sialylation signatures and the clearest responses to enzymatic desialylation. In HD T cell co-cultures, desialylation increased activation, proliferation, cytokine production, and TCE-mediated tumor-cell killing. In primary CLL samples, combined glofitamab and AUS treatment enhanced cytotoxic effector transcriptional programming. These findings support a role for surface sialoglycans in limiting the responsiveness of less-differentiated human T cells to TCE stimulation.

These findings extend current concepts of sialidase-based immunotherapy by providing compartment-specific evidence that desialylation of human T cells directly enhances TCE responsiveness. Prior work has largely focused on tumor- or microenvironment-directed desialylation, with effects reported across tumor, myeloid^14,16,63^, lymphoid^15,22,64^, and stromal compartments^65,66^. Our results build on studies using antibody-sialidase fusion proteins and T cell-targeted sialidase formats, as well as reports that desialylation can enhance T cell activation during antigen-driven^10,21,23^ or anti-CD3 stimulation^67^. In our co-culture experiments, selective desialylation of the T cell compartment was sufficient to augment TCE-induced activation. In the Raji/glofitamab model, Siglec-7/9 blockade did not increase activation, and the sialidase effect persisted after deletion of target-cell CD80/CD86, arguing that neither Siglec-7/9 engagement nor CD28 costimulation accounted for the early activation phenotype under these conditions. These results establish a direct T cell contribution to the sialidase effect without assigning it to a single inhibitory pathway.

We extended the analysis to primary CLL samples to determine whether sialidase treatment alters glofitamab-induced T cell programs in a human disease context where T cell dysfunction is common and responses to CD20-directed TCEs remain suboptimal^34–37^. By combining scRNA-seq with lectin-based CITE-seq, we linked T cell subset identity and treatment-induced transcriptional states with lectin-defined cell-surface glycan signatures. This integrated approach was important because glycosylation cannot be inferred reliably from transcriptomes alone^68^. The transcriptional response to glofitamab and AUS treatment was strongly subset-biased. By normalized DEG burden, activated naïve CD4^+^ and CD8^+^ T cell states showed the highest DEG proportions within their respective lineages, and central memory subsets also underwent substantial remodeling. Yet, activated CD8^+^ Temra cells also showed a sizable absolute number of DEGs, indicating that responsiveness was not strictly ordered by differentiation state. In the less-differentiated subsets, combined treatment increased expression of costimulatory TNF receptor superfamily members, including *TNFRSF4*, *TNFRSF9*, and *TNFRSF18*, while reducing quiescence-associated transcripts such as *BACH2*, *SELL*, and *IL7R* and increasing expression of effector-associated genes, including *TNF*, *GNLY*, and *IFNG*. These changes are consistent with enhanced cytotoxic effector-like transcriptional programming. The induction of *TNFRSF4* is notable because TNFRSF4-expressing CD8^+^ T cells and the TNFSF4-TNFRSF4 axis have been associated with responses to blinatumomab^69,70^, raising the possibility that related costimulatory programs may also be relevant to TCE-sialidase combinations.

This subset bias corresponded to differences in lectin-defined glycan signatures and sialylation-associated gene expression. At baseline, naïve and central memory T cells showed elevated MAL II and SNA binding together with higher expression of *ST3GAL1* and *ST6GAL1*, whereas glofitamab-associated activated naïve T cell states showed lower expression of these sialyltransferases and higher *NEU1* expression. These transcriptional changes indicate activation-associated remodeling of sialylation-related gene expression. Interpretation of the lectin data, however, requires consideration of activation-induced blastogenesis, because larger cells can yield higher total per-cell lectin signals without a corresponding increase in glycan density. After FSC normalization as an approximate correction for cell size, proliferating naïve T cells showed reduced MAL II binding and a weaker trend toward reduced SNA binding. Thus, activation-associated remodeling was most evident in α2,3-linked lectin reactivity. This pattern differs from murine studies emphasizing α2,6-linked remodeling and may reflect species-related regulation of T cell sialoglycans, although variation in activation conditions, timing, and analytical approaches may also contribute^18^. Together, these data associate less-differentiated T cell states with elevated baseline lectin-defined sialylation signatures and suggest that this glycan state may help explain their sensitivity to sialidase treatment, such that exogenous AUS reinforces this intrinsic trajectory and lowers the naïve activation threshold of early differentiated T cells.

We next considered whether specific α2,3-sialylated glycoproteins contribute to the sialoglycan-dependent restraint on TCE responsiveness. CD43 was a strong candidate because it is a heavily O-glycosylated, mucin-like T cell surface protein, carries ST3GAL-dependent α2,3-sialylation, and can limit T cell activation through its bulky, negatively charged ectodomain and effects on receptor organization at the immunological synapse^71,72^. The reported shift from highly sialylated CD43 glycoforms in naïve T cells towards less-sialylated forms during effector differentiation is consistent with the activation-associated decrease in Siglec-7-Fc binding observed here^57^, although this probe reports total Siglec-7 ligands rather than CD43-specific glycoforms^60^. Functionally, CD43 knockout increased activation in response to glofitamab alone and attenuated, but did not abolish, the additional effect of AUS, most clearly in CD4^+^ T cells. Because CD43 deletion removes the entire protein, these experiments cannot distinguish the contribution of its α2,3-linked glycans from effects of the protein scaffold or ectodomain. Moreover, the persistence of AUS-enhanced activation after PMA/ionomycin stimulation suggests that CD43-related effects on immunological synapse organization are unlikely to fully account for the sialidase response. CD43 therefore emerges as a major carrier of α2,3-linked sialoglycans and a partial contributor to AUS responsiveness, suggesting that additional sialylated surface components also contribute to the regulation of TCE-mediated T cell activation. Other heavily sialylated membrane proteins such as CD8α^53,73^, P-selectin glycoprotein ligand-1 (PSGL-1)^74^, and CD44^75,76^ may also contribute, but because neither their expression nor their sialylation is restricted to naïve T cells they are less likely to individually explain the naïve-biased phenotype and were not prioritized here.

Our study has several limitations that affect the generalizability and translational interpretation of the findings. First, the functional experiments were performed in short-term *in vitro* co-cultures and primarily interrogated peripheral T cells. Although enhanced activation was also observed with TCEs directed against solid tumor antigens, these systems did not model chronically stimulated or exhausted T cells within a complex tumor microenvironment. Human-relevant *in vivo* models or advanced *ex vivo* systems that reproduce sustained antigen exposure, stromal interactions, and tissue architecture will therefore be needed to determine how sialidase treatment affects T cell trafficking, tumor infiltration, response durability, and anti-tumor efficacy. Second, the primary CLL analysis was limited to transcriptional and lectin-defined readouts, without direct functional validation. Finally, the mechanisms underlying the stronger response of less-differentiated T cell subsets remain incompletely defined. Further studies will be needed to determine the relative contributions of α2,3- and α2,6-linked sialic acids and of individual sialylated surface molecules beyond CD43.

The preferential sensitivity of naïve and central memory T cells to desialylation may be therapeutically relevant because these populations can provide expandable effector pools, particularly in settings where suboptimal priming can imprint durable T cell dysfunction^77^. The induction of *CXCR3* in naïve T cell states after combined glofitamab and AUS treatment further raises the possibility that desialylation influences migration-associated transcriptional programs, although effects on trafficking, peripheral recruitment, and response durability remain to be established *in vivo*^51,80^. Together with the early clinical evaluation of sialidase-based immunomodulation, including a first-in-human bisialidase-Fc construct (NCT05259696)^79^, our findings support further investigation of sialidase-TCE combinations as a strategy to increase the contribution of less-differentiated T cells to anti-tumor responses.

## Material and methods

### Cell lines

Lenti-X HEK293T were purchased from Takara Bio (RRID: CVCL_4401). Jurkat (ACC282, RRID: CVCL_0065) were purchased from DSMZ, Leibniz Institute. Raji (ATCC CCL-86, RRID: CVCL_0511) and A375 (ATCC CRL-1619; RRID: CVCL_0132), OVCAR8 (human ovarian carcinoma, RRID: CVCL_1629) and SHP-77 (human lung carcinoma, ATCC CRL-2195; RRID: CVCL_2195) were purchased from ATCC. Cell lines were maintained in either DMEM high glucose or RPMI 1640 (Capricorn Scientific) supplemented with 10% fetal bovine serum (FBS), penicillin/streptomycin (100 IU/mL and 100 µg/mL), and 1% (v/v) MEM non-essential amino acids. RPMI 1640 was additionally supplemented with 1% (v/v) sodium pyruvate, 1% (v/v) HEPES, and 0.1% (v/v) 2-mercaptoethanol. Cells were cultured at 37 °C and 5% CO□ and passaged at 70-80% confluency every 2-3 days using trypsin-EDTA in adherent cell lines. Cells were kept in culture for a maximum of 4 weeks.

### Primary chronic lymphocytic leukemia and healthy donor blood cells

Peripheral blood samples from patients diagnosed with CLL were obtained following written informed consent in accordance with the Declaration of Helsinki. The study was approved by the institutional ethics committees of the Universities of Hamburg-Eppendorf, Halle-Wittenberg, and Basel. Detailed clinical characteristics of the patient cohort are provided in **Supplementary Table 1**. Primary human PBMCs were isolated from healthy donor buffy coats (Blutspende SRK Nordwestschweiz) or from patient peripheral blood by density gradient centrifugation using lymphocyte separation medium and SepMate™-50 tubes (STEMCELL Technologies). Cells were cryopreserved in 90% FBS and 10% DMSO and stored in liquid nitrogen until use.

T cells were cultured in complete RPMI 1640 supplemented with 10% FBS, penicillin/streptomycin, L-glutamine, and recombinant human IL-2 (100 IU/mL, Proleukin®), as indicated in the respective assays up to a maximum of 5 days culture time.

### Generation of modified cell lines

Raji CD80/CD86 double-knockout cell line: Low passage Raji cells were electroporated with sgRNA-Cas9 complexes targeting human CD80 and CD86 (**Supplementary Table 2**) using the Maxcyte GT platform with the Expanded T Cell 2 program and PBS. sgRNA and Cas9 nuclease V3 were obtained from IDT. Cells were then expanded for 7 days before sorting for CD80^−^CD86^−^ live cells using an AriaIII sorter (BD Biosciences) and rested for further 7 days.

A375-GFP-CD20/Raji-dTomato-Luc2: For cytotoxicity assay, A375 cells were transduced lentivirally with GFP and CD20 and Raji cells were transduced with dTomato-Luc2. Therefore, lentivirus was produced as previously described^80^. In short, Lenti-X 293T cells were seeded 16 hours before transfection. For each construct, pMD2.G (2.5 μg; RRID: Addgene_12259), pCMVR8.74 (4.7 μg; RRID: Addgene_22036), and the lentiviral transfer plasmid (7.2 μg, constructs listed in **Supplementary Table 3**) were combined in 1.8 mL jetOPTIMUS buffer (Polyplus, Cat. 101000025), followed by addition of 18 μL jetOPTIMUS; complexes were incubated for 10 min and then applied to the cells. Medium was replaced 5-6 h post-transfection, and viral supernatants were harvested 24 h and 48 h after the medium change, pooled and concentrated using a 4× in-house prepared PEG-8000 solution. Concentrated virus was resuspended in 1 mL DMEM, aliquoted, and stored at −80 °C.

For lentiviral transduction, low passage A375 cells were sequentially transduced with CD20 and GFP. After 7 days, stably integrated GFP^+^CD20^+^ cells were sorted with an AriaIII sorter (BD Biosciences) and expanded for further 7 days. Low passage Raji cells were transduced as described and also sorted 7 days after transduction with an AriaIII sorter (BD Biosciences).

### Functional healthy donor T cell assays

For functional assays, cryopreserved human PBMCs were thawed, washed in PBS, and rested for 4 h in complete RPMI prior to assay setup. Naïve pan T cells or total pan T cells were isolated from healthy donor PBMCs using the EasySep™ Human Naïve Pan T Cell Isolation Kit (STEMCELL Technologies, Cat. Nr: 17961) or the EasySep™ Human T Cell Isolation Kit (STEMCELL Technologies, Cat. Nr: 17951), respectively, according to the manufacturer’s instructions. T cell assays were performed in round-bottom 96-well plates using 1.0 × 10^5 T cells per well in a final volume of 200 µL RPMI supplemented with IL-2 (100 IU/mL, Proleukin®).

For activation assays, naïve or total pan T cells were stimulated with ImmunoCult™ CD3/CD28 T Cell Activator (STEMCELL Technologies, Cat. Nr: 10991; 10 µL/mL) and cultured in the presence or absence of 10 U/mL *A. ureafaciens* sialidase (Lectenz Bio, Cat. Nr: GE0701) for a maximum of 5 days prior to downstream analysis at the indicated time points. For proliferation assays, cells were labeled with CellTrace™ Violet (CTV, Thermo Fisher Scientific) prior to stimulation, and proliferation was assessed by dye dilution gating on proliferated CTV^−^ cells.

To assess TCR-independent activation, T cells were stimulated with 0.1 µg/mL PMA and 3 µg/mL ionomycin for 4 h prior to flow-cytometric analysis.

To analyze CD28-independent effects, T cells were stimulated on non-tissue-culture-treated, flat-bottom 96-well plates coated with Ultra-LEAF™ purified anti-CD3 (OKT3; BioLegend, Cat. Nr: 317347; 1 µg/mL) for 24 h at 37 °C, followed by flow-cytometric analysis.

For co-culture assays, adherent target cells were seeded 1 day prior to assay, and suspension target cells were seeded on the day of assay. Pan T cells were added at an effector-to-target (E:T) ratio of 5:1 together with bispecific T cell engagers (glofitamab, blinatumomab, tarlatamab, CD3×EpCAM; Universitätsspital Basel) in the presence or absence of *A. ureafaciens* sialidase (10 U/mL, Lectenz Bio, Cat. Nr: GE0701) or *S. pneumoniae* sialidase (10 U/mL, Lectenz Bio, Cat. Nr: GE0302). For heat-inactivated sialidase controls, sialidase preparations were incubated for 60 min at 80 °C before use.

Where indicated, cells were pre-treated with *A. ureafaciens* sialidase (10 U, Lectenz Bio, Cat. Nr: GE0701) in PBS at 37 °C and 5% CO□ for 30 min, followed by two thorough washes with PBS before co-culture.

To dissect Siglec-dependent effects, co-culture activation assays were performed in the presence of blocking antibodies against Siglec-7 (R&D Systems, Cat. Nr: MAB1138) or Siglec-9 (R&D Systems, Cat. Nr: MAB1139), alongside an isotype-matched IgG2a control (R&D Systems), as indicated.

### Cytotoxicity assay

Cytotoxicity of Raji target cells was assessed using BioTracker NucView® 405 Blue Caspase-3 Dye (Sigma-Aldrich, Cat SCT102). Co-cultures were established as described using Raji-dTomato-Luc2 target cells. After 48 h of co-culture, the fluorogenic caspase-3 substrate was added to each well at a final concentration of 1 µM and incubated for 45 min at 37 °C, 5% CO□. Cells were then washed with FACS buffer, stained with Zombie NIR viability dye, fixed, and analyzed by flow cytometry. Untreated cells and heat-shocked cells (60 °C, 20 min) were included as negative and positive controls, respectively, to define gating thresholds.

Live-cell killing was monitored using an IncuCyte® S3 live-cell imaging system under standard culture conditions. Images were acquired every 4 h. Target cell killing of A375-CD20□GFP□ cells was quantified based on loss of GFP signal over time combined with AI-based confluence analysis provided by the instrument software. GFP signal was normalized to the time point = 0 h.

### Flow cytometry and cell sorting

Cells were pelleted by centrifugation, washed twice with PBS, and stained with fluorochrome-conjugated antibodies and/or lectins in 50 µL PBS per well (full panel in **Supplementary Table 4**). Viability dyes (Zombie™ Fixable Viability Kits) and Fc receptor-blocking reagents were included where indicated. Surface staining was performed for 20 min at 4 °C, followed by washing in ice-cold FACS buffer (PBS, 0.5 mM EDTA, 2% FBS, and 0.1% sodium azide (NaN□)), fixation in eBioscience™ IC Fixation Buffer, and resuspension in FACS buffer for acquisition.

For intracellular cytokine staining, 1× Monensin (BioLegend) was added for the final 4 h of culture prior to flow-cytometric analysis. For degranulation assays, anti-CD107a-APC-H7 (BD Biosciences, Cat. Nr: 561343; 20 ng/mL final) was included during the final 4 h of culture.

For intranuclear or intracellular staining, cells were fixed and permeabilized using Fixation/Perm diluent (Invitrogen) according to the manufacturer’s instructions, and subsequently stained with antibody cocktails in 1× permeabilization buffer (Invitrogen). Full antibody panel is listed in **Supplementary Table 4**. Samples were acquired on Cytek® Aurora, CytoFLEX S, or BD LSRFortessa™ cytometers and analyzed using FlowJo v10.10. Cell sorting was performed on BD SORP FACSAria or CytoFLEX SRT cell sorters.

### Cytokine measurements

Cytokines in culture supernatants were quantified using the LEGENDplex™ Human CD8/NK multiplex panel (BioLegend, Cat. Nr: 741187) according to the manufacturer’s protocol after 24 hours of co-cultures. Samples were acquired on a Cytoflex S and analyzed using the LEGENDplex™ analysis software.

### Conjugation of lectins with oligonucleotides

Lectins conjugated to DNA oligonucleotides, each carrying a unique barcode sequence for precise identification and a capture sequence compatible with the 10x Genomics Next GEM kit, were produced as recently described^81^. Unconjugated lectins, including *Aleuria aurantia* lectin (AAL), *Maclura pomifera* agglutinin (MPA), concanavalin A (ConA), *Datura stramonium* agglutinin (DSA), wheat germ agglutinin (WGA), peanut agglutinin (PNA), *Maackia amurensis* lectin I (MAA I), and *Erythrina cristagalli* agglutinin (ECA), as well as biotinylated *Maackia amurensis* lectin II (MAL II) and *Sambucus nigra* agglutinin conjugated to FITC (SNA-FITC), were obtained from Vector Laboratories (Newark, CA, USA). The table of lectins and their unique barcode identifiers is listed in **Supplementary Table 5**.

### T cell engager activation assay with PBMC from patients with CLL

Cryopreserved peripheral blood mononuclear cells (PBMCs) from patients with CLL (*n* = 5) were rapidly thawed, washed, and rested for 4 h in complete RPMI 1640 medium. Cells were then cultured overnight at 37 °C in a humidified 5% CO□ incubator under the following conditions: (i) CLL (ii) CLL + AUS (iii) CLL + glofitamab, or (iv) glofitamab + AUS. Glofitamab concentration was 1 pM and enzyme concentration was 10 U/mL AUS. Heat-inactivated AUS (HI-AUS, 10 U/mL) was applied to control for buffer-mediated effects in (i), (iii). To enrich for CD19□ immune cells, 10% of the total cell suspension was combined with the flow-through fraction from CD19-positive selection, performed using the EasySep™ Human CD19 Positive Selection Kit II (STEMCELL Technologies, Cat Nr: 17854) according to the manufacturer’s instructions. The resulting CD19-depleted fraction was used for subsequent flow cytometric sorting.

### Surface Staining for CITE-seq and Flow Cytometric Sorting

Cells were washed and resuspended in PBS containing 1% BSA, Ca²□/Mg²□, and Human Fc Receptor Binding Inhibitor (Thermo Fisher Scientific). After incubation for 20□min at 4□°C, cells were washed and further incubated on ice for 30□min with DNA-barcoded lectins, MAL II-biotin and SNA-FITC, each at 0.5□µg/mL (0.05□µg per 10□□cells). In parallel, surface antibody CITE-seq staining was performed using oligonucleotide-conjugated antibodies against CD45RA (clone HI100, Biolegend TotalSeqB), CD45RO (clone UCHL1, Biolegend TotalSeqB), and CCR7 (clone G043H7, Biolegend TotalSeqB), together with conventional fluorescence surface markers: CD3 PE (clone UCHT1), CD45 BV711 (clone HI30), CD4 APC (clone SK3), CD8 PE-Cy7 (clone SK1), and CD19 BV650 (clone HIB19). TotalSeq B (BioLegend) anti-human Hashtag 1 (0251), Hashtag 2 (0252), Hashtag 3 (0253), Hashtag 5 (0255), and Hashtag 6 (0256) antibodies were also added at this stage according to the manufacturer’s instructions. A secondary labeling step was carried out using streptavidin-oligonucleotide (Biolegend, TotalSeq-B0975) and anti-FITC-oligonucleotide (Biolegend, TotalSeq B-0988, clone: FIT-22) conjugates. Following incubation, cells were washed twice and resuspended in staining buffer containing DAPI (5 µg/mL; Thermo Fisher Scientific) for live/dead discrimination.

Fluorescence-activated cell sorting (FACS) was performed on a BD-FACSAria-III. Single, live (DAPI-negative), CD45□, and CD19□□cells were purified into cold PBS containing 20%□FBS (Sigma-Aldrich), spiking in 10% of CD19+ B cells. Sorted cells were merged, pooling the donors per experimental conditions and were immediately processed for single-cell immune profiling using the 10x□Genomics Chromium platform.

### Single-cell glycan and RNA sequencing

Single-cell capture, cDNA and library preparation were performed at the Genomics Facility of the University of Basel, with a Single-Cell 3′ v4 Reagent Kit with Feature Barcode technology for Cell Surface Protein (10x Genomics). Oligonucleotide-conjugated lectins were used to produce antibody derived tag (ADT) libraries. Sequencing was performed on the AVITI instrument in four runs resulting in 28nt-long and 90nt-long paired-end reads, respectively.

cDNA reads were aligned to ‘hg38’ genome using Ensembl 110 gene models with the STARsolo tool (v2.7.9a) with default parameter values except the following parameters: soloUMIlen=12, soloBarcodeReadLength=1, outFilterType=BySJout, outFilterMultimapNmax=10, outSAMmultNmax=1, soloType=CB_UMI_Simple, outFilterScoreMin=30, soloCBmatchWLtype=1MM_multi_Nbase_pseudocounts, soloUMIfiltering=MultiGeneUMI_CR, soloUMIdedup=1MM_CR, soloCellFilter=None. ADT libraries were mapped to the reference ADT dictionary in the same way.

Merged samples from different donors were demultiplexed based on present single nucleotide polymorphisms (SNPs). For this purpose tools cellsnp-lite^82^ and vireo^83^ were applied. As the input BAM files with mapped reads and the SNP database HapMap 3.3 (https://www.sanger.ac.uk/resources/downloads/human/hapmap3.html) were used. As a result, cells were either assigned to a particular genotype of individual donors or not assigned to any genotype.

Further analysis was performed using R (v4.4.1). Cell filtering was done based only on the analysis of the gene expression, not ADT abundance. The package DropletUtils (v1.22.0) was applied to detect empty droplets and define the ambient profile.

Cells were considered as high-quality cells if they had at least 600 detected genes, which is the threshold derived from the distribution of the number of detected genes across cells, and assigned to a specific donor forming a data set of 86980 cells. The function scDblFinder from the package scDblFinder (v1.18.0) pointed at 6451 doublets, which were removed from the further analysis resulting in 80529 cells.

Normalization, integration across donors and conditions and clustering were performed using the package Seurat (v4.4.1). The resulting Seurat object was converted to a SingleCellExperiment (v1.26.0) object and the rest of the analysis was performed using multiple Bioconductor (v3.19) packages. The data set was subjected to the cell-type annotation using the Bioconductor package SingleR (v2.6.0) and samples from Human Primary Cell Atlas provided by the Bioconductor package celldex (v1.14.0) as the reference.

Clusters of cells assigned to T cells or NK cells were extracted (Fig. 3B) and re-clustered to purify the T-cell population and reveal different T-cell states, which were defined by exploring the expression of specific marker genes (**Fig. S9A**).

For performing differential expression analysis, pseudo-bulk samples were created by summing up counts per gene across all cells belonging to a particular cluster, treatment, and donor. The function ‘filterByExpr’ excluded lowly expressed genes from the analysis. Combinations containing at least 20 cells underwent the analysis with edgeR (4.2.2). Genes were considered as differentially expressed if false discovery rate (FDR)<0.05. Gene set enrichment analysis was performed using the tool camera from the edgeR package and the Gene Ontology gene set from MSigDB (v2023.2). The analysis of changes in the abundance of clusters assigned to specific T-cell states across treatments was also performed with edgeR. Z-scores for the normalized expression of pseudo-bulk samples were used to generate heatmaps, i.e. “z-score[log2(cpm)]”. For the heatmap visualization of the ADT expression, normalized abundance of a glycan was averaged across all cells belonging to a particular cluster, treatment, and donor and z-scores were calculated, i.e. “z-score[logcounts]”. Lower and upper limits of z-scores were set to −2 and 2, respectively.

### CRISPR-Cas9-mediated CD43 knockout in primary T cells

CD43 knockout was generated in freshly isolated human pan T cells using Cas9 ribonucleoprotein (RNP) electroporation. Negative control crRNA #1 (IDT, Cat. Nr: 1072544) and Alt-R™ tracrRNA (IDT, Cat. Nr: 1072533) were annealed according to the manufacturer’s protocol. Non-targeting gRNA or CD43-targeting sgRNA (5′-GGCTCGCTAGTAGAGACCAA-3′; 150 pmol) were complexed with recombinant Cas9 Nuclease V3 protein (IDT, Cat. Nr: 1081058; 60 pmol). RNPs were electroporated into T cells using the MaxCyte GT platform with the Expanded T Cell 2 program in PBS. Cells were immediately transferred into pre-warmed complete PrimeXV medium (Irvine Scientific) for 4 h, and subsequently cultured in PrimeXV supplemented with IL-7 (10 ng/mL), IL-15 (10 ng/mL), and 1% penicillin-streptomycin. Three days after electroporation, CD43□ T cells were sorted on a FACSAria III cell sorter (BD Biosciences) and used for co-culture assays as described above. For lectin-staining controls, T cells were isolated and activated for 2 days with ImmunoCult™ CD2/CD3/CD28 T Cell Activator (STEMCELL Technologies, Cat. Nr: 10990; 25 µL/mL).

### Statistics and visualization

The statistical analysis and graph preparation were performed using the software package Prism version 11.0.0 (GraphPad Software, La Jolla, CA). Statistical tests are specified in the corresponding figure legends, where P values > 0.05 were considered not significant, and P values < 0.05 were considered significant. *P < 0.05, **P < 0.01, ***P < 0.001, and ****P < 0.0001. Flow cytometry data was analysed using FlowJo version 10.10.0 followed by Prism version 11. R version 4.4.1 and BioRender.com were used for visualizations.

### AI-assisted manuscript review

Before submission, the manuscript was reviewed using Large Langugag models (ChatGPT 5.6, Occam’s Pen). All conceptual content and conclusions were developed independently by the authors and take full responsibility for all content in the submitted manuscript.

## Acknowledgements

We thank all of the members of the Cancer Immunology and Cancer Immunotherapy Laboratory of the Department of Biomedicine for helpful discussions. We are grateful to all of the patients who provided material. We thank Dominic Schwarz, Dario Venetz and Christian Klein (Roche Glycart, RICZ, Switzerland) for sharing EPCAM-TCB. We also thank Irene Fusi (Cancer Immunology, Department of Biomedicine, University of Basel) for contributing to the manuscript review. Furthermore, we want to thank the Flow Cytometry and Genomics Core Facilities of the Department of Biomedicine for their support. We also thank Dominik Burri for the discussion on experimental design and Florian Geier for pre-processing CITE-seq data. Calculations were performed at sciCORE (http://scicore.unibas.ch/) scientific computing center at University of Basel.

## Contributions

JN and HL planned the project and experimental design. JN, DH, AnZ, and MS performed the *in vitro* experiments. AB and MS processed and analyzed the RNA-sequencing data and performed the data deposition. JN, DH, MS, AB, LG, and HL interpreted the results. RM and PA provided critical reagents, including custom lectin-oligonucleotide conjugates and advised on lectin-based glycan profiling. CS and MB provided clinical samples, collected clinical data, and obtained ethical approval. JN, LG, and HL wrote the manuscript, A. Zippelius provided important input in formulating and revising the manuscript. All authors reviewed and approved the final version.

## Data, code, and materials availability

All data needed to evaluate the conclusions in the paper are present in the paper or the Supplementary Materials. Because of data security guidelines of the University of Basel, raw sequencing data are under restricted access. Researchers can apply for access by contacting the Data Access Committee (DAC) at. EpCAM TCE is available from Dario Venetz under a material transfer agreement (MTA) with Roche, Switzerland. Lectin-oligonucleotides are available from Pablo Argüeso under a MTA with Tufts University, USA.

All other materials are commercially available as described in Materials and Methods.

