## Supplemental Data for "α2,3-sialylation on human naïve T cells restrains bispecific engager-mediated anti-tumor immunity"

### Supplementary material

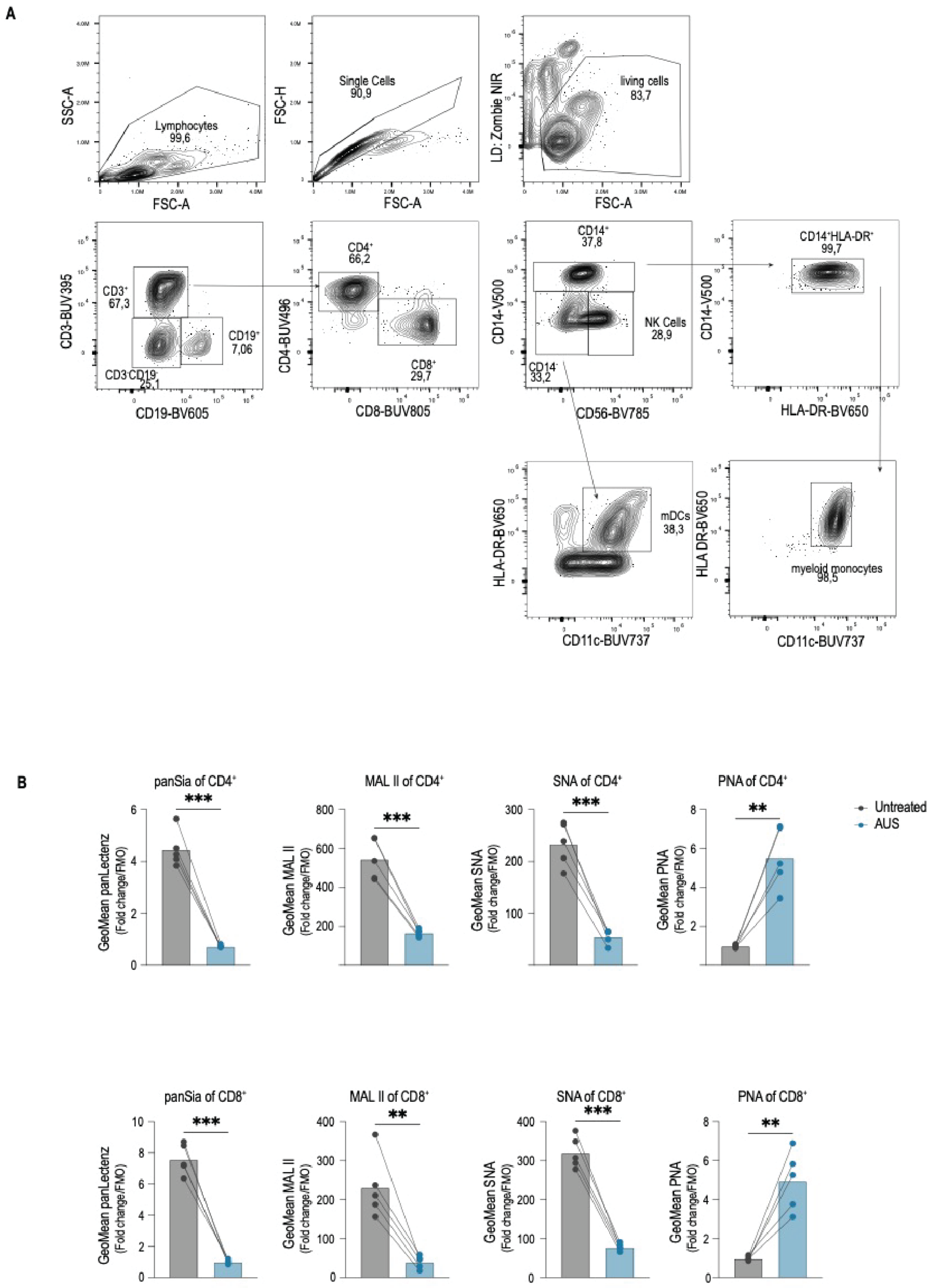


**Fig. S1: Flow-cytometric assessment of human T cell sialylation and desialylation**

**A)** Representative contour plots showing the gating strategy used to identify PBMC subpopulations. T cells were defined as Singlet/Live/CD3$^{+}$, B cells as Singlet/Live/CD19$^{+}$, NK cells as Singlet/Live/CD3$^{-}$CD19$^{-}$/CD56$^{+}$, DCs as Singlet/Live/CD3$^{-}$CD19$^{-}$/CD14$^{-}$/HLA-DR$^{+}$CD11c$^{+}$, Monocytes as Singlet/Live/CD3$^{-}$CD19$^{-}$/CD14$^{+}$/HLA-DR$^{+}$CD11c$^{+}$. **B)** GeoMean fluorescence intensity of lectin binding normalized to the corresponding FMO control in CD4$^{+}$ and CD8$^{+}$ T cells after treatment ± 10 U/mL AUS. Lectins shown include Pan lectenz, MAL II, SNA, and PNA ($n$ = 5 donors). Data are shown as means, samples from the same donor are connected by lines. Statistical significance was assessed by two-tailed paired Student's $t$ test. **$p < 0.01$, ***$p < 0.001$, ****$p < 0.0001$. Abbreviations: AUS, *Arthrobacter ureafaciens* sialidase; DC, dendritic cells; FMO, fluorescence-minus-one; GeoMean, geometric mean; MAL II, *Maackia amurensis* lectin II; NK, natural killer cells; PBMCs, peripheral blood mononuclear cells; PNA, peanut agglutinin; SNA, *Sambucus nigra* agglutinin.

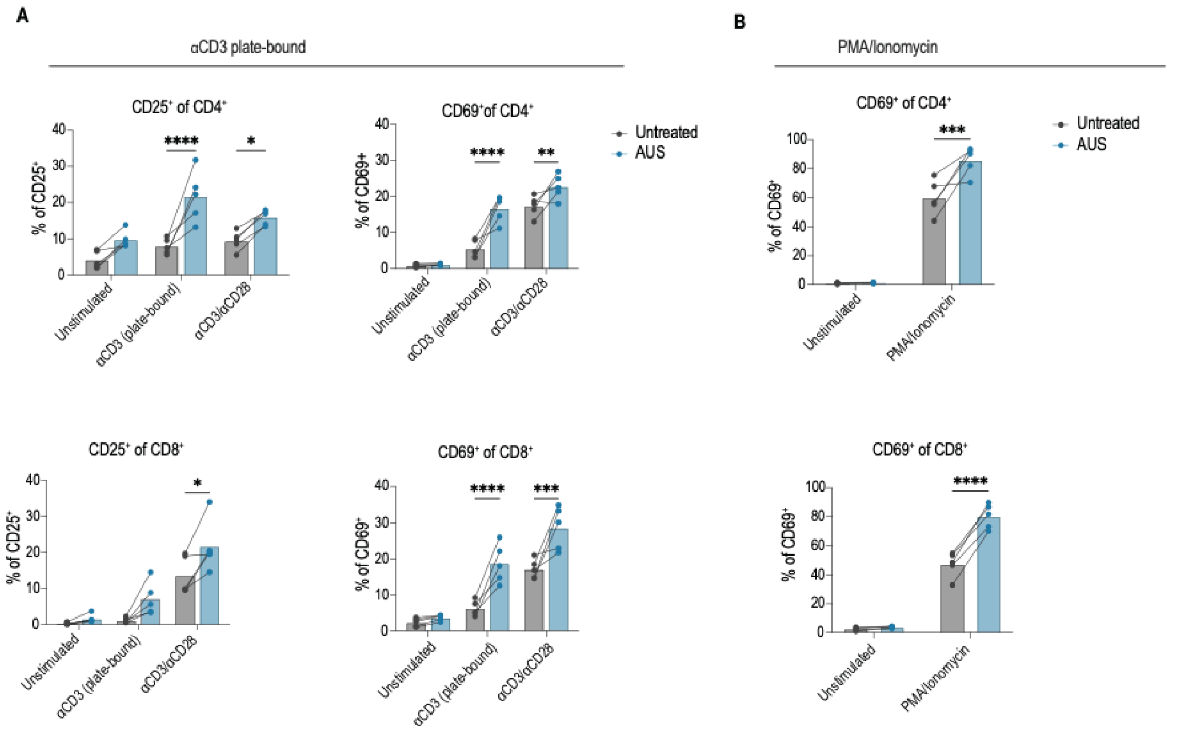


**Fig. S2: CD28-independent and PMA/ionomycin stimulation controls for AUS-enhanced T cell activation**

A) Frequencies of activated CD25+ and CD69+ CD4+ and CD8+ T cells after stimulation with plate-bound αCD3 (1 µg/mL) or αCD3/αCD28 (10 µl/mL) for 24 hours ± 10 U/mL AUS (n = 5 donors). B) Frequencies of CD69+ CD4+ and CD8+ T cells after stimulation with (PMA; 0.1 µg/mL) and ionomycin (3 µg/mL) for 4 hours ± 10 U/mL AUS (n = 5 donors). Data are shown as mean values, samples from the same donor are connected by lines. Statistical significance was assessed by two-way ANOVA with Holm-Sidak correction. *p < 0.05, **p < 0.01, ***p < 0.001, ****p < 0.0001. Abbreviations: AUS, Arthrobacter ureafaciens sialidase; PMA, phorbol 12-myristate 13-acetate.

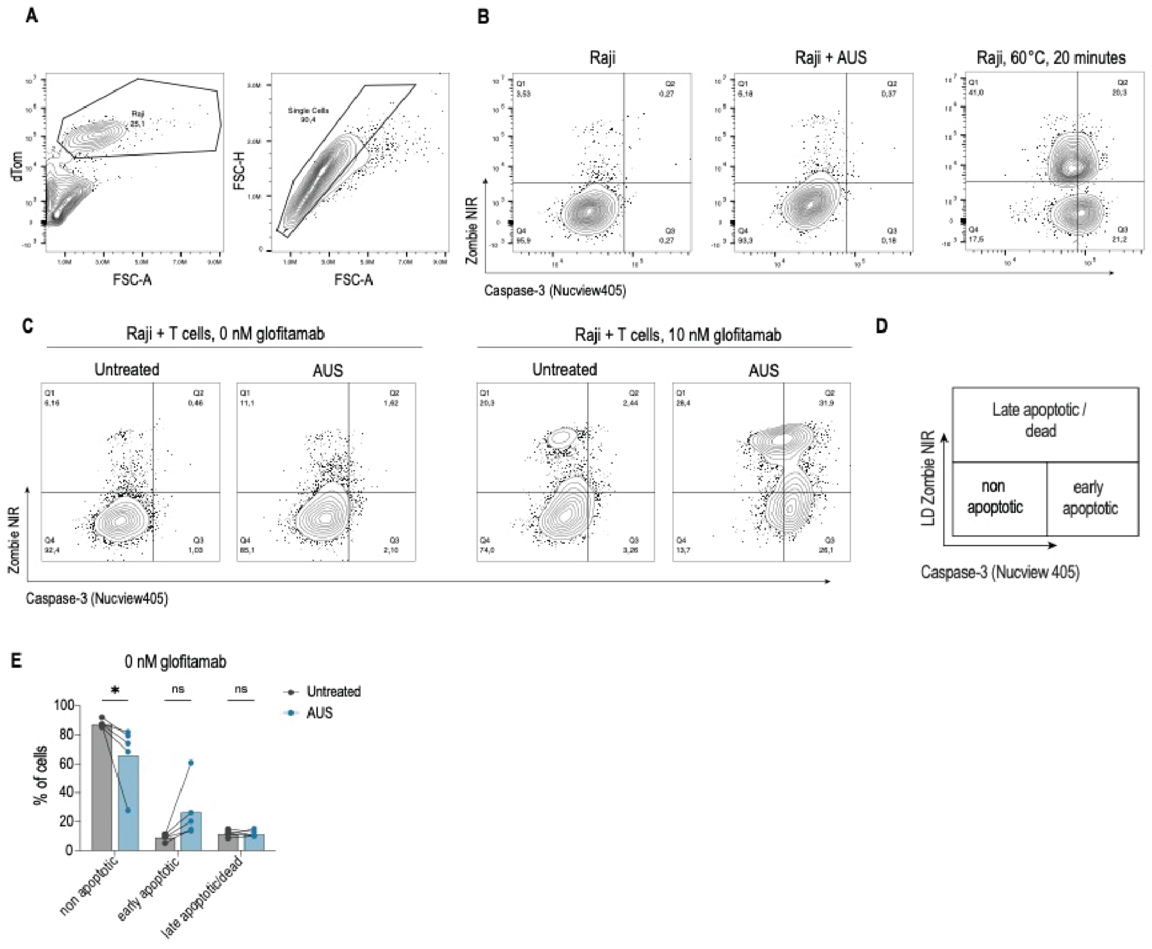


**Fig. S3: Controls for glofitamab-mediated target-cell killing assays**

**A, B)** Flow cytometric assessment of caspase-3 activity in Raji target cells using BioTracker NucView 405. Control conditions included untreated Raji cells, AUS-treated Raji cells, and heat-treated dead-cell controls (heated for 20 minutes at 60 °C). **C)** Representative contour plots of caspase-3 and Zombie NIR staining after co-culture with T cells in the absence or presence of 10 pM glofitamab and ± 10 U/mL AUS **D)** Gating strategy used to define non-apoptotic, early apoptotic, and late apoptotic/dead target cells. Non-apoptotic cells were defined as Zombie NIR$^{-}$caspase-3$^{-}$, early apoptotic cells as Zombie NIR$^{-}$caspase-3$^{+}$, and late apoptotic/dead cells as Zombie NIR$^{+}$caspase-3$^{-/+}$. **E)** Frequencies of caspase-3$^{+}$ Raji target cells after 48 hour co-culture with T cells ± 10 U/mL AUS in the absence of glofitamab; $n$ = 5 donors. Statistical significance was assessed by one-way ANOVA with Holm-Sidak correction. *$p < 0.05$, **$p < 0.01$, ***$p < 0.001$, ****$p < 0.0001$. Abbreviations: AUS, *Arthrobacter ureafaciens* sialidase; Zombie NIR, near infrared fixable viability dye.

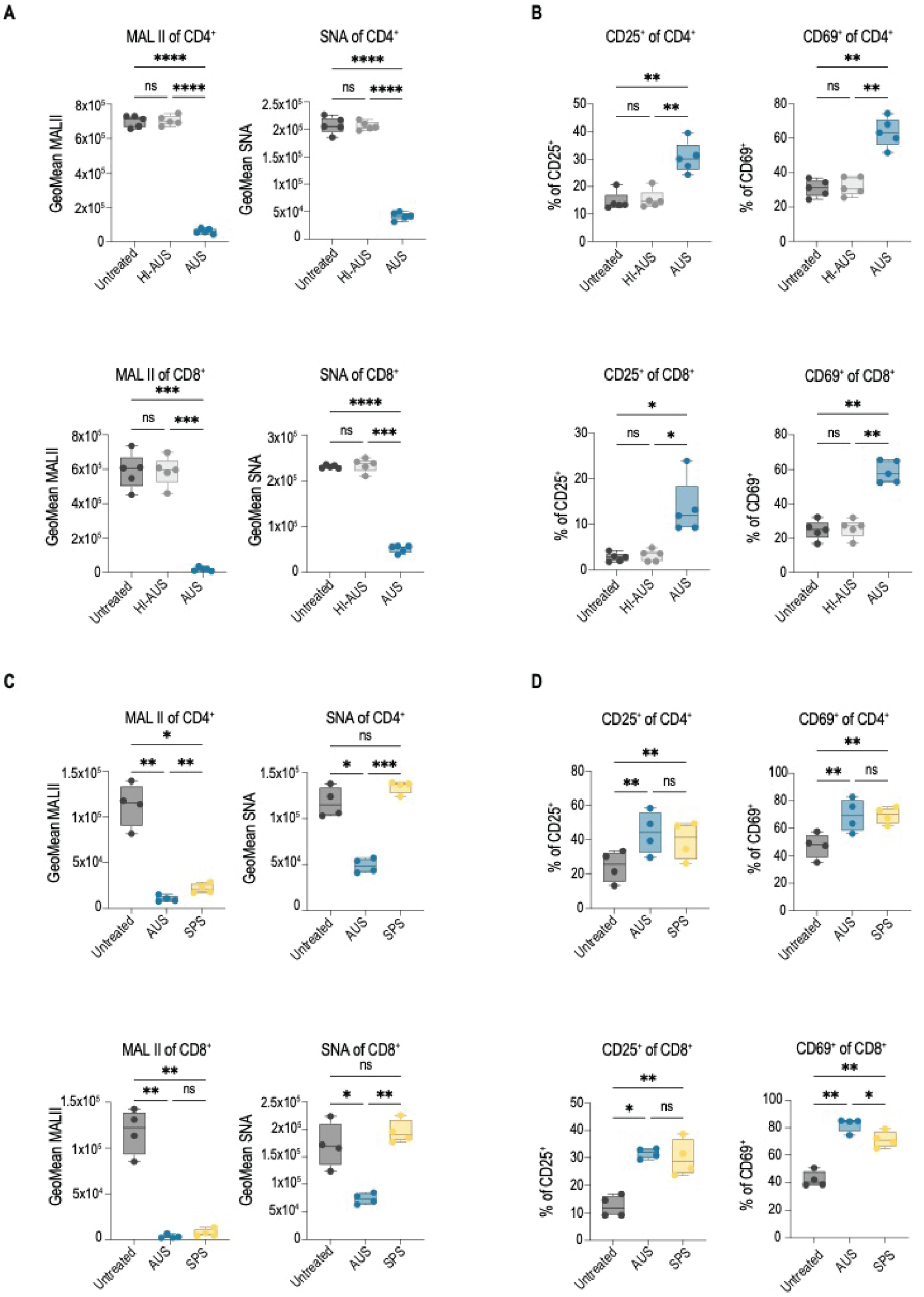


**Fig. S4: Heat-inactivated and α2,3-specific sialidase controls in TCE co-cultures**

**A)** MAL II and SNA binding on CD4$^+$ and CD8$^+$ T cells after Raji/glofitamab co-culture with untreated control, HI-AUS, or enzymatically active AUS; *n* = 5 donors. **B)** Frequencies of CD25$^+$ and CD69$^+$ CD4$^+$ and CD8$^+$ T cells under the conditions shown in **A**. **C)** MAL II and SNA binding on CD4$^+$ and CD8$^+$ T cells after Raji/glofitamab co-culture with untreated control, AUS, or SPS; *n* = 4 donors. **D)** Frequencies of CD25$^+$ and CD69$^+$ CD4$^+$ and CD8$^+$ T cells under the conditions shown in **C**. Co-cultures were stimulated with 0.1 pM glofitamab. Enzymes were used at 10 U/mL. Statistical significance was assessed by one-way ANOVA with Tukey correction. *$p < 0.05$, **$p < 0.01$, ***$p < 0.001$, ****$p < 0.0001$. Abbreviations: AUS, *Arthrobacter ureafaciens* sialidase; HI-AUS, heat-inactivated AUS; MAL II, *Maackia amurensis* lectin II; SNA, *Sambucus nigra* agglutinin; SPS, *Streptococcus pneumoniae* α2,3-sialidase; TCE, T cell engager.

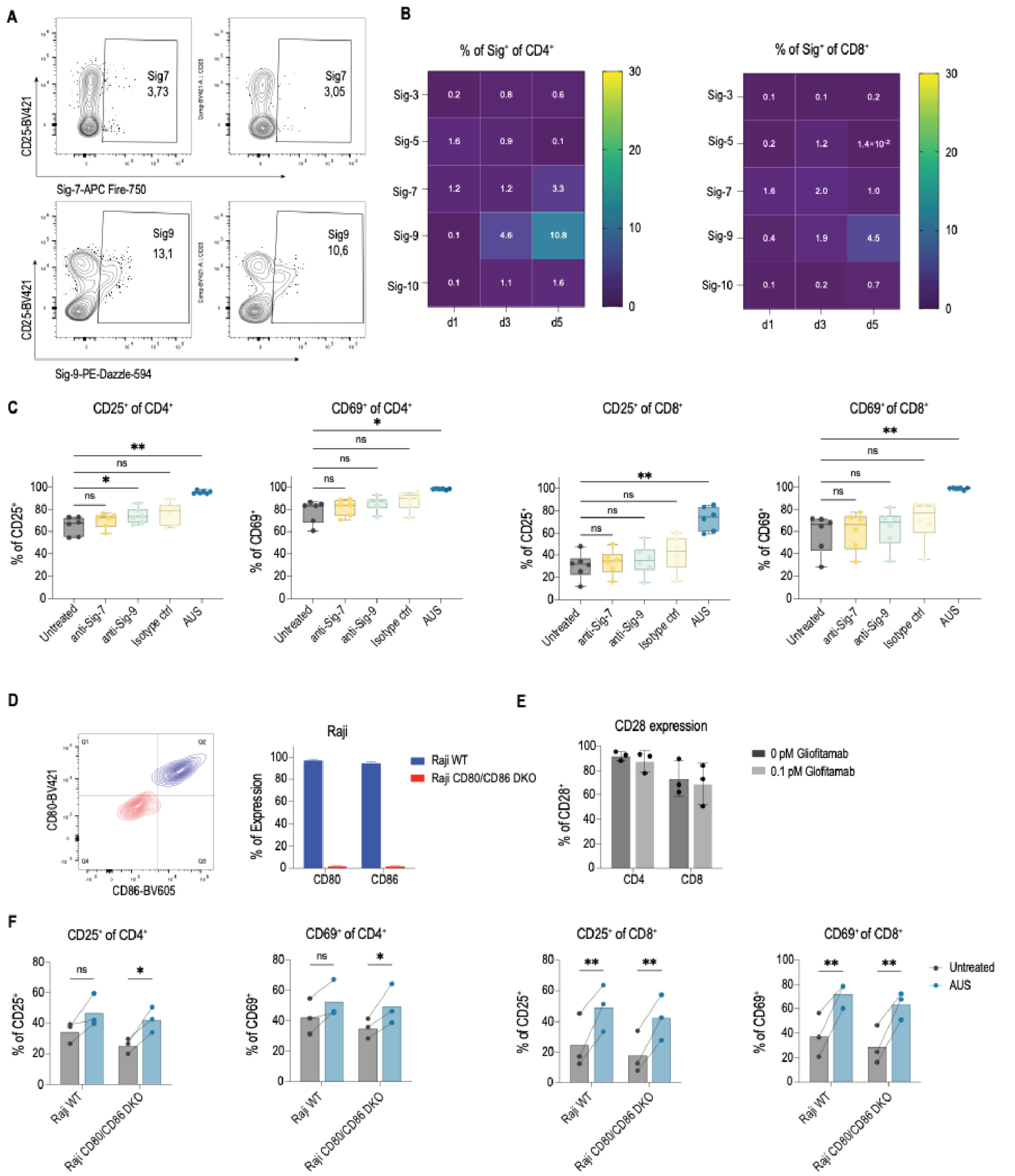


**Fig. S5: Siglec-7/9- and CD28-mediated mechanism controls for AUS-enhanced T cell activation**

**A)** Representative contour plots of Siglec-7 and Siglec-9 expression on CD4$^{+}$ and CD8$^{+}$ T cells after 5 days of Raji/glofitamab co-culture. **B)** Frequencies of Siglec-3$^{+}$, Siglec-5$^{+}$, Siglec-7$^{+}$, Siglec-9$^{+}$, and Siglec-10$^{+}$ CD4$^{+}$ and CD8$^{+}$ T cells on days 1, 3, and 5 after stimulation with Raji cells and 0.1 pM glofitamab; $n$ = 5 donors. **C)** Frequencies of CD25$^{+}$ and CD69$^{+}$ CD4$^{+}$ and CD8$^{+}$ T cells after 24-hour Raji/glofitamab co-culture in the presence of 10 µg/mL Siglec-7 and Siglec-9 blocking antibodies, IgG2a isotype control, or 10 U/mL AUS; $n$ = 5 donors. **D)** Representative contour plots and frequencies of CD80 and CD86 expression on wild-type Raji cells and Raji CD80/CD86 DKO cells. **E)** CD28 expression on CD4$^{+}$ and CD8$^{+}$ T cells after 24-hour co-culture with Raji cells ± 0.1 pM glofitamab; $n$ = 3 donors. **F)** Frequencies of CD25$^{+}$ and CD69$^{+}$ CD4$^{+}$ and CD8$^{+}$ T cells after 24-hour co-culture with wild-type or CD80/CD86 DKO Raji cells and 0.1 pM glofitamab ± 10 U/mL AUS; $n$ = 3 donors. Statistical significance was assessed by one-way ANOVA with Tukey correction for **C** and by two-way ANOVA with Holm–Sidak correction for **F**. *$p < 0.05$, **$p < 0.01$. Abbreviations: AUS, Arthrobacter ureafaciens sialidase; DKO, double knockout; Siglec, sialic acid-binding immunoglobulin-like lectin; WT, wild type.

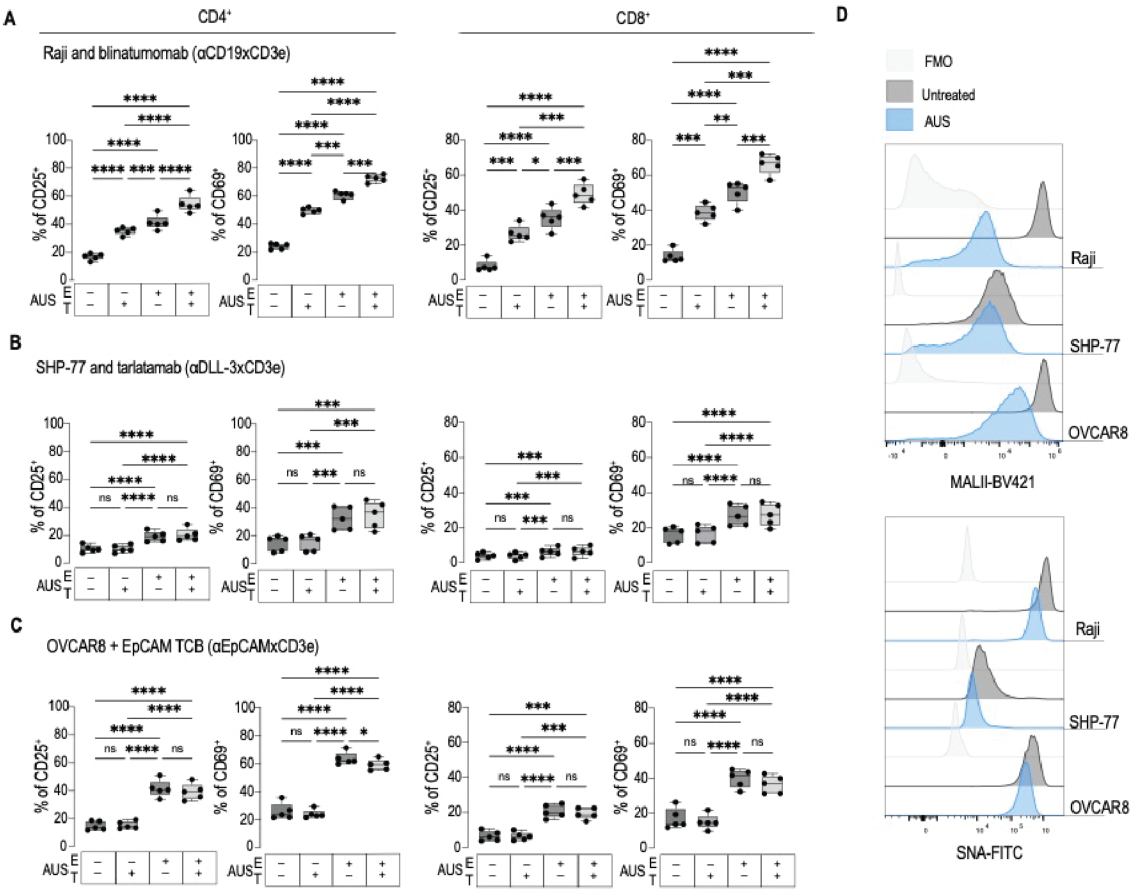


**Fig. S6: T cell-directed desialylation enhances activation across additional TCE models**

**A-C)** Frequencies of CD25+ and CD69+ CD4+ and CD8+ T cells after co-culture with target cells and the indicated TCEs following compartment-specific desialylation of effector T cells and/or target cells. Effector T cells and target cells were pretreated separately with AUS, washed to remove residual enzyme, and then combined for co-culture. "+" indicates AUS treatment; "–" indicates untreated control. **A)** CD19+ Raji cells stimulated with 1 pM blinatumomab. **B)** DLL3+ SHP-77 cells stimulated with 10 pM tarlatamab. **C)** EpCAM+ OVCAR-8 cells stimulated with 0.5 pM CD3×EpCAM TCB. **D)** Representative histograms of MAL II and SNA binding on the target-cell lines used in **A–C** before and after AUS treatment. Statistical significance was assessed by one-way ANOVA with Tukey correction; $n = 5$ donors. $^{*}p < 0.05$, $^{**}p < 0.01$, $^{***}p < 0.001$, $^{****}p < 0.0001$. Abbreviations: AUS, *Arthrobacter ureafaciens* sialidase; MAL II, *Maackia amurensis* lectin II; SNA, *Sambucus nigr*a agglutinin; TCB, T cell bispecific antibody; TCE, T cell engager.

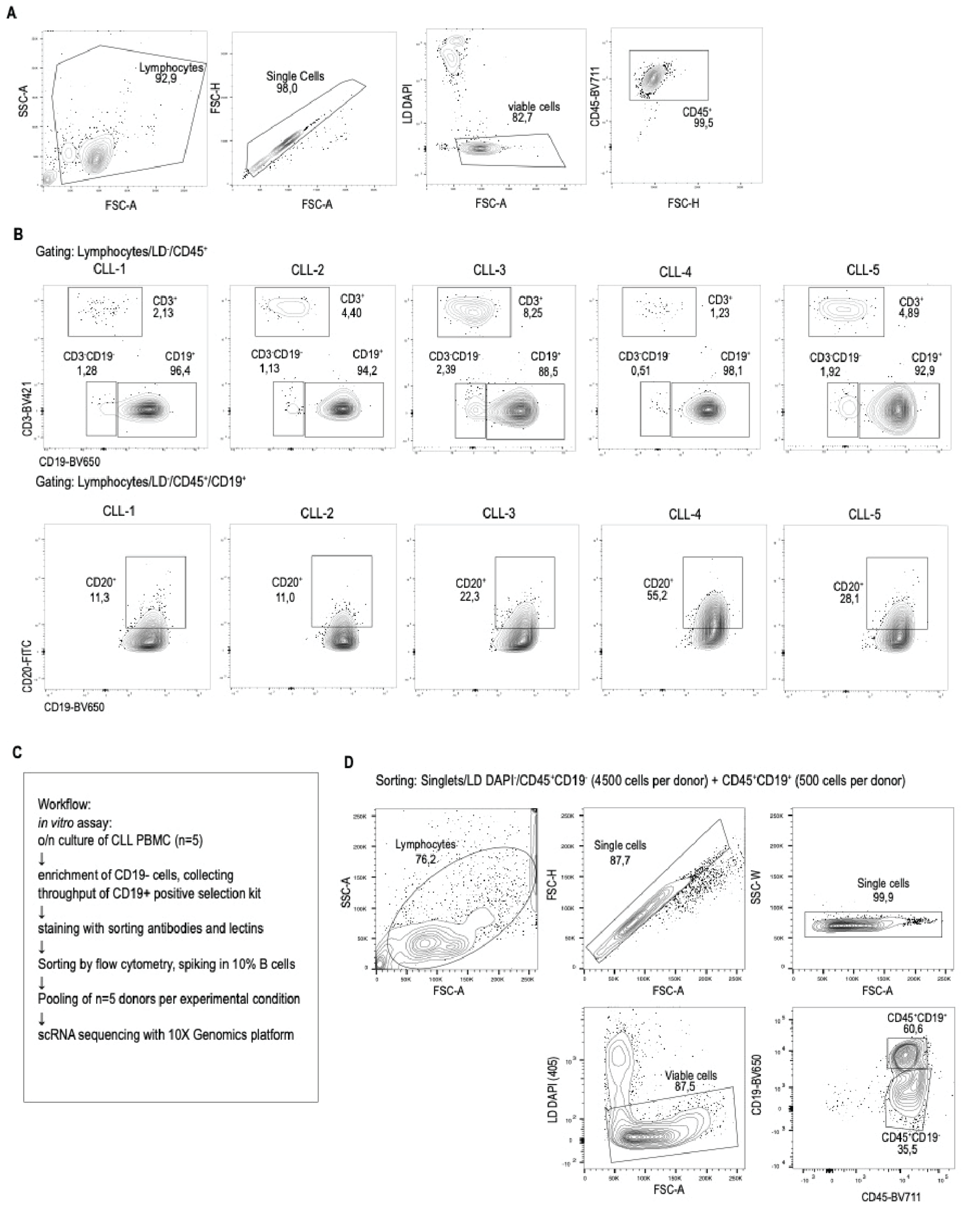


**Fig. S7: CLL sample composition and sorting workflow for single-cell glycan profiling**

**A)** Workflow of the CLL *ex vivo* culture and single-cell profiling experiment. PBMCs from patients with CLL were cultured overnight under the indicated treatment conditions, followed by CD19-based enrichment of non-B immune cells, antibody and lectin staining, flow-cytometric sorting of live CD45$^+$ cells with a defined CD19$^+$ fraction included, pooling by treatment condition, and downstream scRNA-seq with lectin-based CITE-seq. **B)** Flow-cytometric characterization of CLL PBMC samples. Representative gating strategy for lymphocytes, single cells, viable cells, and CD45$^+$ cells. **C)** Top panel: Patient-level plots showing frequencies of CD45$^+$CD3$^+$ T cells, CD45$^+$CD19$^+$ B cells, and CD45$^+$CD3$^-$CD19$^-$ immune cells across the 5 CLL donors. Bottom panel: CD20 expression within the CD45$^+$CD19$^+$ B cell compartment. **D)** Representative flow-sorting strategy after CD19-based enrichment. Single viable CD45$^+$ cells were sorted as CD45$^+$CD19$^-$ immune cells and

$CD45^{+}CD19^{+}$ B cells, with 4500 $CD45^{+}CD19^{-}$ cells and 500 $CD45^{+}CD19^{+}$ cells collected per donor to generate an approximate 90:10 $CD19^{-}$ to $CD19^{+}$ input for downstream single-cell profiling. Abbreviations: CITE-seq, cellular indexing of transcriptomes and epitopes by sequencing; CLL, chronic lymphocytic leukemia; PBMCs, peripheral blood mononuclear cells; scRNA-seq, single-cell RNA sequencing.

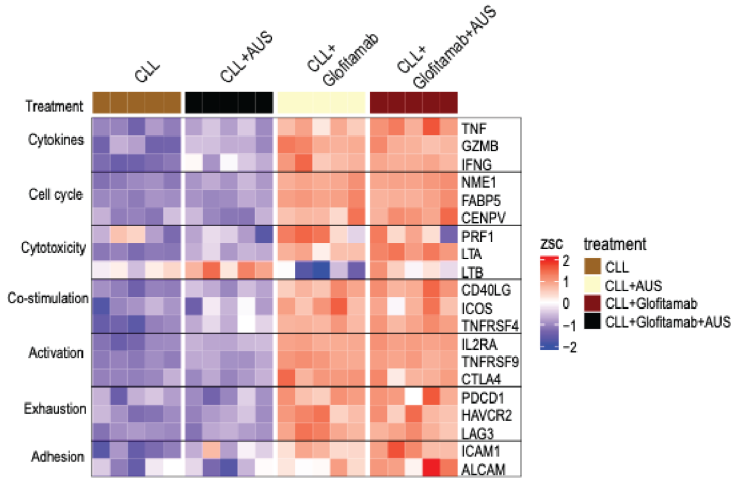


**Fig. S8: Glofitamab and sialidase induce activation and effector transcriptional programs CLL T cells**

Heatmap showing expression of selected T cell gene sets associated with activation, cytotoxic effector function, proliferation, and related immune functions across the 4 *ex vivo* CLL culture conditions: CLL, CLL + AUS, CLL + glofitamab, and CLL + glofitamab + AUS. Expression values are shown as z scores of normalized pseudo-bulk gene expression, calculated as z-score[log2(cpm)]. Z scores were capped at −2 and 2 for visualization. Abbreviations: AUS, *Arthrobacter ureafaciens* sialidase; CLL, chronic lymphocytic leukemia; cpm, counts per million; zsc, z scores.

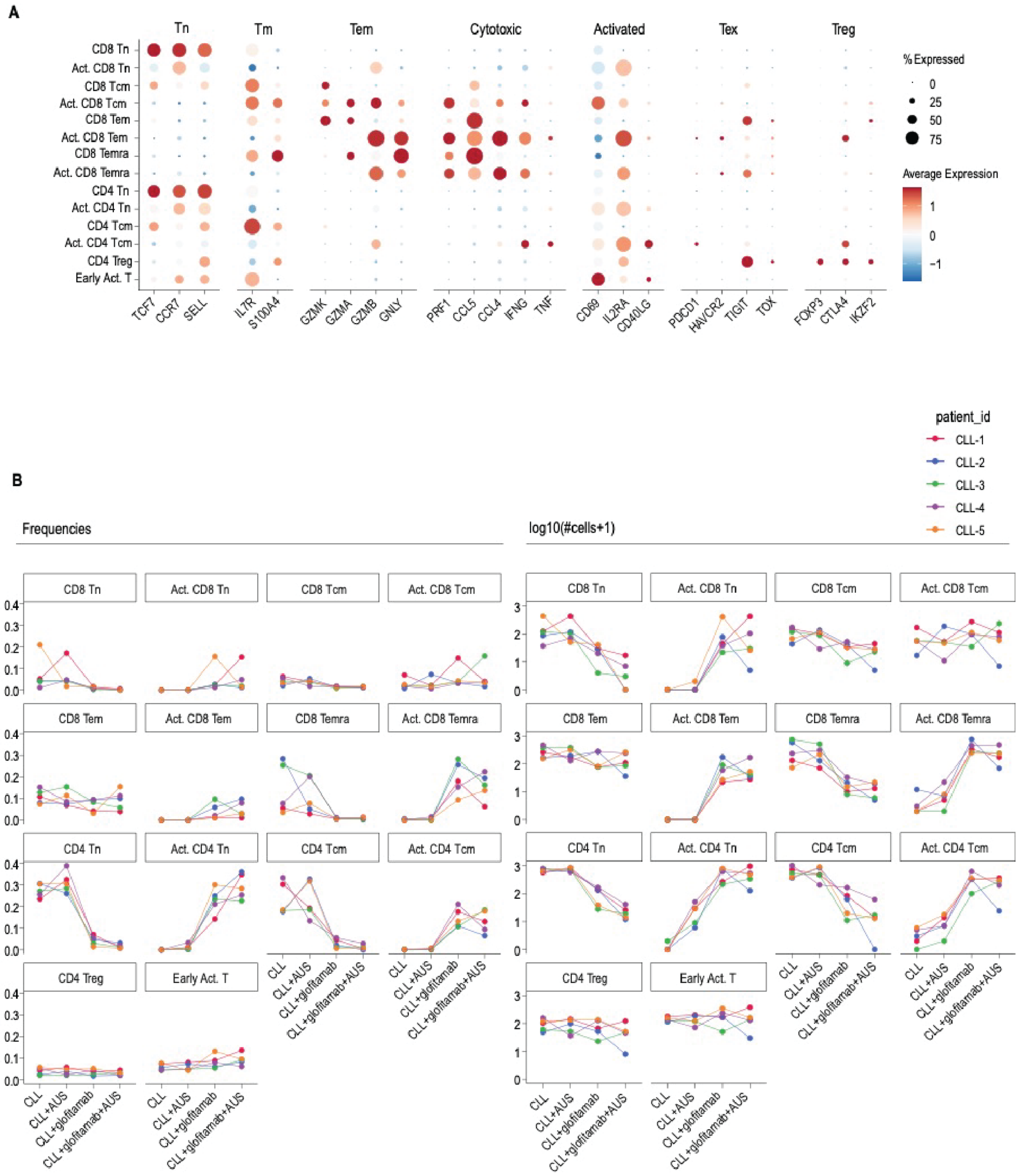


**Fig. S9: Annotation and treatment distribution of CLL T cell subclusters**

**A)** Dot plot showing expression of canonical marker genes used to annotate T cell subclusters. Dot size indicates the percentage of cells expressing each gene, and color indicates z-scored average expression within each subcluster. Marker groups are shown above the plot to indicate naïve, memory, effector, cytotoxic, activated, exhaustion-associated, and regulatory T cell features. **B)** Donor-level frequencies and cell numbers for each annotated T cell subcluster across the 4 treatment conditions: CLL, CLL + AUS, CLL + glofitamab, CLL + glofitamab + AUS. Left, frequency of each subset among T cells; right, log10-transformed cell counts per subset. Lines connect matched samples from the same CLL donor across treatment conditions. Abbreviations: AUS, *Arthrobacter ureafaciens* sialidase; CLL, chronic lymphocytic leukemia; Tcm, central memory T cells; Tem, effector memory T cells; Temra, terminally differentiated effector memory T cells re-expressing CD45RA; Tn, naïve T cells; Treg, regulatory T cells.

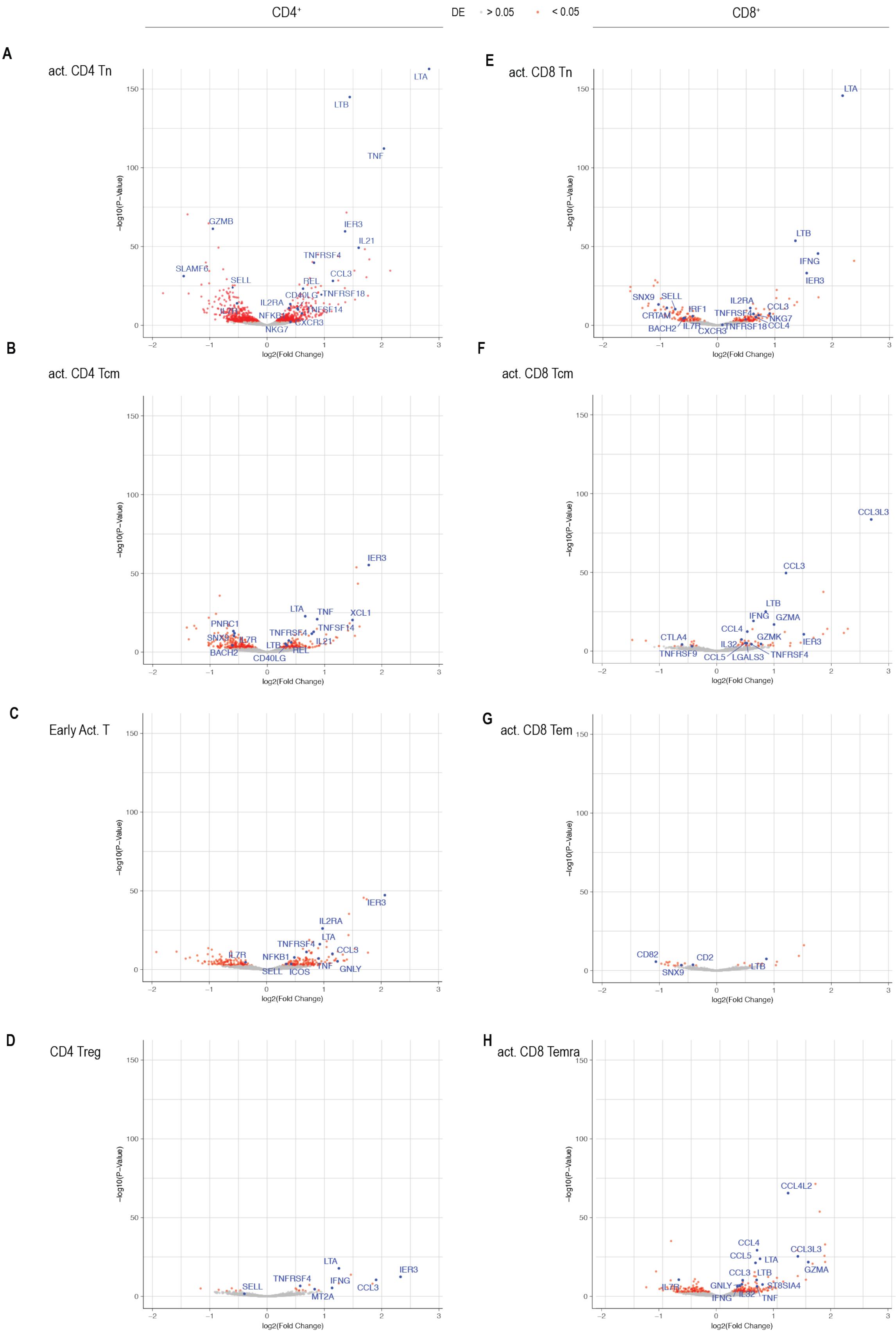


**Fig. S10: Subset-specific differential gene expression after combined glofitamab and AUS treatment**

**A-H)** Volcano plots showing DEGs within annotated CD4$^{+}$ and CD8$^{+}$ T cell subpopulations comparing CLL + glofitamab + AUS with CLL + glofitamab alone. Positive log2 fold changes indicate higher expression after combined glofitamab and AUS treatment; negative values indicate higher expression after glofitamab alone. Genes passing absolute log2 fold change > 0.3 and Benjamini–Hochberg FDR-adjusted p <

0.05 are highlighted; selected genes are annotated. Abbreviations: AUS, *Arthrobacter ureafaciens* sialidase; CLL, chronic lymphocytic leukemia; DEG, differentially expressed gene; FDR, false discovery rate; Tcm, central memory T cells; Tem, effector memory T cells; Temra, terminally differentiated effector memory T cells re-expressing CD45RA; Tn, naïve T cells; Treg, regulatory T cells.

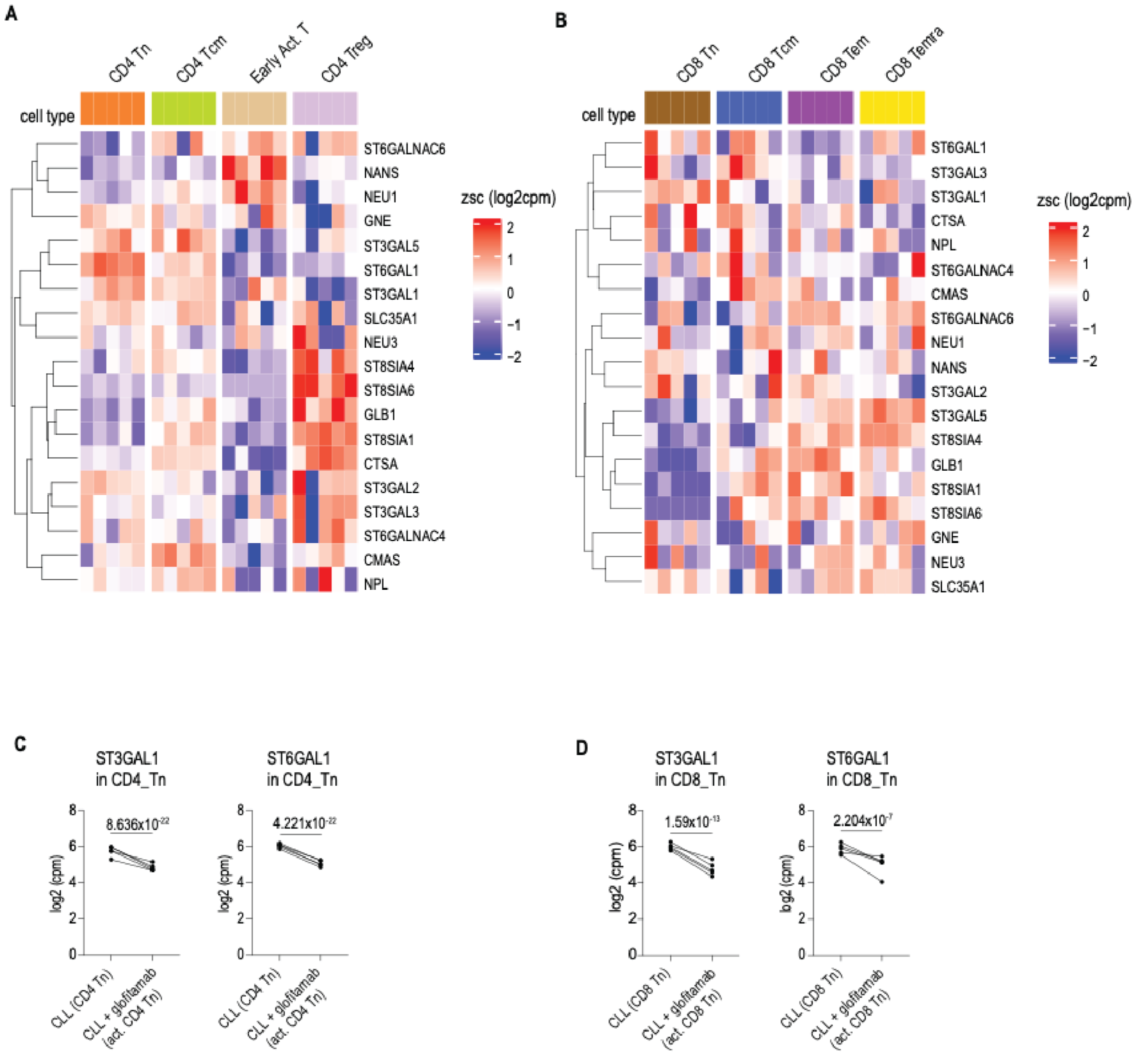


**Fig. S11: Sialylation-associated gene expression in CLL T cell subsets**

**A,B)** Heatmaps showing RNA expression of selected sialylation-associated genes in CD4$^{+}$ **(A)** and CD8$^{+}$ **(B)** T cell subsets from untreated CLL PBMC cultures. Values are shown as row-wise z scores of normalized log2[cpm], capped at -2 and 2 for visualization. **C,D)** Paired donor-level expression of ST3GAL1 and ST6GAL1 in resting naïve CD4$^{+}$ T cells versus activated naïve CD4$^{+}$ T cells after glofitamab stimulation **(C)** and resting naïve CD8$^{+}$ T cells versus activated naïve CD8$^{+}$ T cells after glofitamab stimulation **(D)**. Values are shown as log2(cpm), with paired samples from the same donor connected by lines. Statistical values indicate Benjamini–Hochberg FDR-adjusted *p* values. Abbreviations: AUS, *Arthrobacter ureafaciens* sialidase; CLL, chronic lymphocytic leukemia; cpm, counts per million; FDR, false discovery rate; Tcm, central memory T cells; Tem, effector memory T cells; Temra, terminally differentiated effector memory T cells re-expressing CD45RA; Tn, naïve T cells; Treg, regulatory T cells.

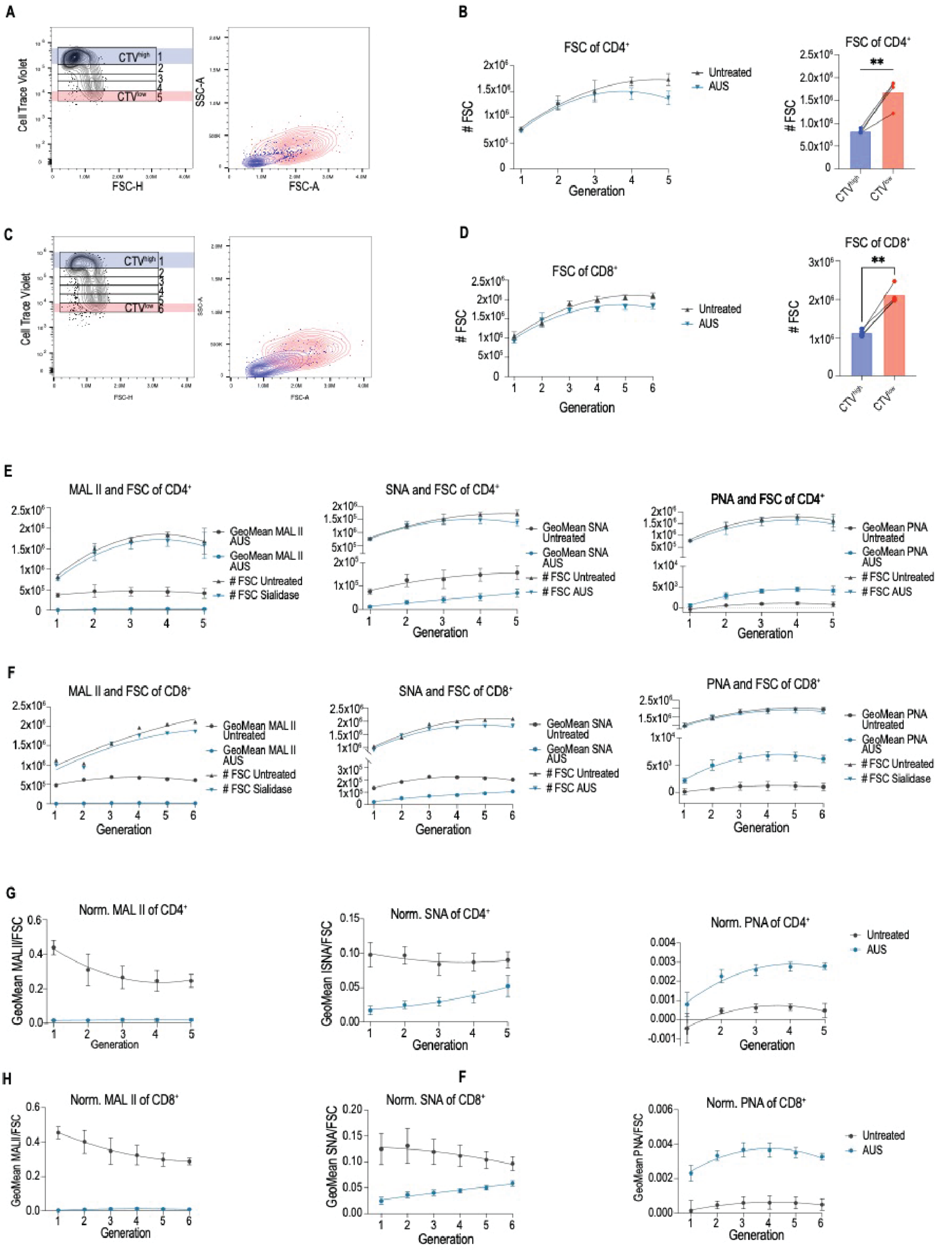


**Fig. S12: Cell size-adjusted lectin analysis during naïve T cell activation**

**A,C)** Representative flow-cytometric plots of naïve CD4+ **(A)** and CD8+ **(C)** T cells after 5-day activation with αCD3/αCD28. CTV dilution was used to define proliferative generations. Undivided cells in generation 1 were defined as CTV$^{high}$, and cells in the latest proliferative generation were defined as CTV$^{low}$. Corresponding FSC-A and SSC-A plots show the cell-size and granularity profile of CTV$^{high}$ and CTVI$^{ow}$ cells. **B,D)** FSC across CTV-defined generations of naïve CD4+ **(B)** and CD8+ **(D)** T cells activated in the presence or absence of 10 U/ml AUS. Bar plots compare FSC between CTV$^{high}$ and CTV$^{low}$ populations. **E,F)** GeoMean fluorescence intensity of MAL II, SNA, and PNA binding plotted together with FSC across CTV-defined generations of naïve $CD4^+$ **(E)** and $CD8^+$ **(F)** T cells activated in the presence or absence of AUS. Curves show nonlinear regression fits; points indicate mean values from $n = 4$ donors. **G**,**H)** Lectin binding from **E** and **F** normalized to FSC to account for activation-associated increases in cell size. Values are shown as GeoMean/FSC across CTV-defined generations for naïve $CD4^+$ **(G)** and $CD8^+$ **(H)** T cells. Curves show nonlinear regression fits; points indicate mean values from $n = 4$ donors. Data are shown for $n = 4$ healthy donors. Where bar plots compare $CTV^{high}$ and $CTV^{low}$ populations, paired samples from the same donor are connected by lines. Statistical significance was assessed using paired two-tailed Student's t test. $^{**}p < 0.01$. Abbreviations: AUS, *Arthrobacter ureafaciens* sialidase; CTV, CellTrace Violet; FSC, forward scatter; GeoMean, geometric mean fluorescence intensity; MAL II, *Maackia amurensis* lectin II; PNA, peanut agglutinin; SNA, *Sambucus nigra* agglutinin; SSC, side scatter.

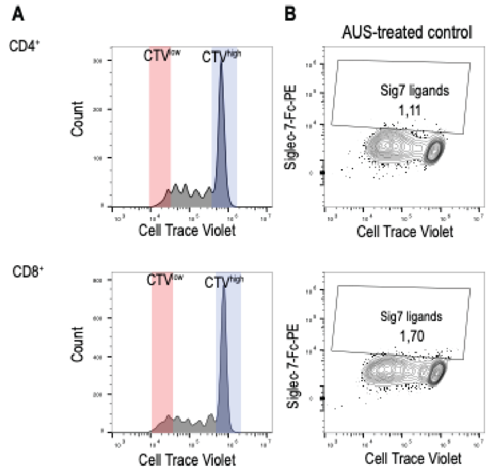


**Fig. S13: Gating controls for Siglec-7-Fc ligand detection**

**A)** Representative histograms of CTV dilution in naïve CD4⁺ and CD8⁺ T cells after 5-day activation with αCD3/αCD28. Undivided CTV$^{high}$ and proliferated CTV$^{low}$ populations were used for analysis of Siglec-7-Fc binding. **B)** Representative contour plots showing Siglec-7-Fc binding plotted against CTV intensity in AUS-treated naïve CD4$^{+}$ and CD8$^{+}$ T cells. AUS-treated cells served as a desialylated control for threshold setting. Abbreviations: AUS, *Arthrobacter ureafaciens* sialidase; CTV, CellTrace Violet.

### Supplementary Tables

| CLL | Age | Sex | Treated at time point of sampling | Comorbidities |
|---|---|---|---|---|
| 1 | 55 | m | untreated for CLL, but chemotherapy for sarcoma | sarcoma |
| 2 | 62 | f | untreated | no |
| 3 | 39 | f | untreated | no |
| 4 | 62 | f | untreated | no |
| 5 | 78 | f | untreated | no |

**Supplementary Table 1: Characteristics of patients with chronic lymphocytic leukemia**

| Target | Sequence | Cat. Nr | Supplier |
|---|---|---|---|
| negative control | na | 1072544 | IDT |
| CD80 | GAAGTGGCAACGCTGTCCTG | | Genescript |
| CD86 | TGTCCGAATCAAAACTTGTG | | Genescript |
| CD43 | GGCTCGCTAGTAGAGACCAA | | IDT |

**Supplementary Table 2: single-guide RNAs for CRISPR/Cas9 knockout**

| Plasmid | Name | Source | Identifier |
|---|---|---|---|
| dtomLuc | pcDNA3.1(+)/Luc2=tdT | Addgene, was a gift from Christopher Contag | RRID:Addgene_32904 |
| CD20 | pLV-CD20 | inhouse | CD20: codon optimized and ordered as fragment from Genscript, uniprot P11836 |
| GFP | pLV-GFP | inhouse | enhanced GFP, ordered via Genscript |

**Supplementary Table 3: Plasmids for lentiviral transduction**

| Name | Color/Label | Clone | Provider | Cat. # |
|---|---|---|---|---|
| Anti-human CCR7 | AF647 | G043H7 | BioLegend, Inc. | 353218 |
| Anti-human CD11c | BUV737 | B-ly6 | BD | 741827 |
| Anti-human CD137 (4-1BB) | BV 786 | 4B4-1 | BD | 741000 |
| Anti-human CD14 | V500 | M5E2 | BD | 561391 |
| Anti-human CD19 | PerCP Cy5.5 | SJ25C1 | BioLegend, Inc. | 363016 |
| Anti-human CD19 | BV605 | SJ25C1 | BioLegend, Inc. | 363024 |
| Anti-human CD2 | PE | RPA-2.10 | Thermo Fisher Scientific Inc. | 12-0029-42 |
| Anti-human CD2 | FITC | S5.2 | BD | 347593 |
| Anti-human CD25 | BV421 | BC96 | BioLegend, Inc. | 302630 |
| Anti-human CD25 | PE-Cy5 | BC96 | BioLegend, Inc. | 302607 |
| Anti-human CD3 | FITC | HIT3a | BioLegend, Inc. | 300306 |
| Anti-human CD3 | BUV 395 | SK7 | BD | 564001 |

| | | | | |
|---|---|---|---|---|
| Anti-human CD4 | APC | SK3 | Thermo Fisher Scientific Inc. | 17-0047-42 |
| Anti-human CD4 | BUV 496 | SK3 | BD | 612936 |
| Anti-human CD43 | FITC | MEM-59 | BioLegend, Inc. | 315203 |
| Anti-human CD43 | APC | 1G10 | BioLegend, Inc. | 560198 |
| Anti-human CD45RA | Pe-Cy7 | HI100 | Thermo Fisher Scientific Inc. | 9025-0458-120 |
| Anti-human CD56 | BV785 | 5.1H11 | BioLegend, Inc. | 362550 |
| Anti-human CD69 | PE-Cy7 | FN50 | BD | 557745 |
| Anti-human CD69 | BV421 | FN50 | BD | 562884 |
| Anti-human CD8a | BV605 | RPA-T8 | BioLegend, Inc. | 301040 |
| Anti-human CD8a | BUV 805 | RPA-T8 | BD | 568334 |
| Anti-human HLA-DR | BV 650 | L243 | BioLegend, Inc. | 307650 |
| Anti-human PD-1 | BV421 | EH12.2H7 | BioLegend, Inc. | 329920 |
| Anti-human Siglec-3 | BV785 | WM53 | BioLegend, Inc. | 303428 |
| Anti-human Siglec-5 | PE | #07 | SinoBiological | 11798-MM07-P |
| Anti-human Siglec-7 | APC fire 750 | 6-434 | BioLegend, Inc. | 339208 |
| Anti-human Siglec-9 | PE-Dazzle | K8 | BioLegend, Inc. | 351516 |
| Anti-human Siglec-10 | PE | 5G6 | BioLegend, Inc. | 347604 |
| Anti-human Tim-3 | PE | F38-2E2 | BioLegend, Inc. | 345006 |
| Siglec-7 Fc | na | na | R&D | 1138-SL |
| anti-human IgG Fc Antibody | PE | M1310G05 | Biolegend | 410708 |
| Lectin PNA | PE | / | GeneTex | GTX01509 |
| Lectin SNA | FITC | / | Vector Laboratories, Inc. | FL-1301 |
| Lectin MAL II | Biotin | / | Vector laboratories | B-1265-1 |
| Streptavidin | BV421 | 500 | BD | 563259 |
| Streptavidin | PE | 500 | BioLegend, Inc. | 405204 |

**Supplementary Table 4: Antibodies and lectins for flow cytometry staining**

| **Lectin/ Antibody** | **Oligo** | **clone** | **Cat. Nr.** | **Name** | **Vendor** | **Remarks** |
|---|---|---|---|---|---|---|
| MPA (*Maclura pomifera* agglutinin) | GAAGAAGCGTTATTC | na | na | na | na | |
| PNA (peanut agglutinin) | GCATTGCGTCAGGCT | na | na | na | na | |
| ECA (*Erythrina cristagalli* agglutinin) | TTAGGTGTACACGTT | na | na | na | na | |
| AAL (*Aleuria aurantia* lectin) | AAGGCAGACGGTGCA | na | na | na | na | |
| MAA-I (*Maackia* | CGAGGTACATCTTGT | na | na | na | na | |

| | | | | | | |
|---|---|---|---|---|---|---|
| *amurensis* lectin I) | | | | | | |
| ConA (concanavalin A) | AGAGCTAGGATCGGA | na | na | na | na | |
| DSA, DSL (*Datura stramonium* agglutinin/lectin) | GGACGCAACTTAAGA | na | na | na | na | |
| WGA (wheat germ agglutinin) | CACTCCTTGACAGGT | na | na | na | na | |
| Streptavidin | TGCCAGCCCTTTGTA | na | 405369 | TotalSeq™-B0975 | Biolegend | 2nd for MALII biot. |
| anti-FITC | TTTGTGTTGTGGTAC | FIT-22 | 408315 | TotalSeq™-B0988 | Biolegend | 2nd for SNA FITC |
| CD45RA | TCAATCCTTCCGCTT | HI100 | 304161 | TotalSeq™-B0063 | Biolegend | |
| CCR7 | AGTTCAGTCAACCGA | G043H7 | 353249 | TotalSeq™-B0148 | Biolegend | |
| CD45RO | CTCCGAATCATGTTG | UCHL1 | 304257 | TotalSeq™-B0087 | Biolegend | |
| Hashtag 1 | GTCAACTCTTTAGCG | LNH-94; 2M2 | 394631 | TotalSeq™-B0251 | Biolegend | 1 |
| Hashtag 2 | TGATGGCCTATTGGG | LNH-94 | 394633 | TotalSeq™-B0252 | Biolegend | 2 |
| Hashtag 3 | TTCCGCCTCTCTTTG | LNH-94 | 394635 | TotalSeq™-B0253 | Biolegend | 3 |
| Hashtag 4 | AAGTATCGTTTCGCA | LNH-94 | 394639 | TotalSeq™-B0255 | Biolegend | 5 |
| Hashtag 5 | GGTTGCCAGATGTCA | LNH-94 | 394641 | TotalSeq™-B0256 | Biolegend | 6 |

**Supplementary Table 5: Antibodies and lectins for cellular indexing of transcriptomes and epitopes by sequencing (CITE-seq)**

| **Category** | **Name** | **Source** | **Identifier** |
|---|---|---|---|
| Critical reagent | ALT-R crRNA | Integrated DNA Technologies (IDT) | Sequence specific - see Supplementary Table 2 |
| Critical reagent | ALT-R tracrRNA | Integrated DNA Technologies (IDT) | 1072532 |
| Critical reagent | Alt-R Cas9 Electroporation enhancer | Integrated DNA Technologies (IDT) | 10007805 |
| Critical reagent | Foxp3 / Transcription Factor Fixation/Permeabilization kit | eBioscience | 00-5521-00 |
| Critical reagent | IC Fixation Buffer | Thermo Fisher Scientific Inc. | 00-8222-49 |

| | | | |
|---|---|---|---|
| Critical reagent | CellTrace CFSE Cell Proliferation Kit | Thermo Fisher Scientific Inc. | C34570 |
| Critical reagent | EasySep Human Naive Pan T Cell Isolation kit | STEMCELL Technologies | 17961 |
| Critical reagent | EasySep Human T Cell Isolation Kit | STEMCELL Technologies | 17951 |
| Critical reagent | ImmunoCult Human CD3/CD28 T Cell Activator | STEMCELL Technologies | 10991 |
| Critical reagent | LEGENDplex Human CD8/NK Panel (13-plex) | BioLegend, Inc. | 741187 |
| Critical reagent | P3 Primary Cell 96-well Nucleofector Kit | Lonza | V4SP-3096 |
| Critical reagent | Pan sialidase | Lectenz Bio, Inc. | GE0701 |
| Critical reagent | α2,3 sialidase | Lectenz Bio, Inc. | GE0302 |
| Critical reagent | Penicillin-streptomycin | Merck KGaA | P4333-100 mL |
| Critical reagent | Permeabilization Buffer (10X) | Thermo Fisher Scientific Inc. | 00-8333-56 |
| Critical reagent | Phorbol 12-myristate 13-acetate (PMA) | Merck KGaA | P8139-1MG |
| Critical reagent | Precision Count Beads | BioLegend, Inc. | 424902 |
| Critical reagent | RBC Lysis Buffer | Thermo Fisher Scientific Inc. | 00-4333-57 |
| Critical reagent | Recombinant human IL-2, Proleukin | SteriMax Inc. | DIN02130181 |
| Critical reagent | Trypan Blue | Merck KGaA | T8154-100 mL |
| Critical reagent | Trypsin (0.05%)-EDTA | LifeTechnologies | 25300054 |
| Critical reagent | DNase I, Type IV | Merck KGaA | D5025 |
| Critical reagent | Fixation/Permeabilization Concentrate | Thermo Fisher Scientific Inc. | 00-5123-43 |
| Critical reagent | Fixation/Permeabilization Diluent | Thermo Fisher Scientific Inc. | 00-5223-56 |
| Critical reagent | EDTA disodium salt | Thermo Fisher Scientific Inc. | 03690-100 mL |
| Critical reagent | 2-Mercaptoethanol | Thermo Fisher Scientific Inc. | 31350-010 |
| Critical reagent | Accutase | Innovative Cell Technologies, Inc. | ICTAT104 |
| Critical reagent | Ambion Nuclease free water | Thermo Fisher Scientific Inc. | AM9938 |
| Critical reagent | Anti-Hu Fc Receptor Binding Inhibitor | Thermo Fisher Scientific Inc. | 14916173 |
| Dye | Zombie UV Fixable Viability Kit | BioLegend, Inc. | 423108 |
| Dye | Zombie Aqua Fixable Viability Kit | BioLegend, Inc. | 423102 |
| Dye | Zombie NIR Fixable Viability Kit | BioLegend, Inc. | 423106 |

| | | | |
|---|---|---|---|
| Dye | CellTrace Violet Cell Proliferation Kit | Thermo Fisher Scientific Inc. | C34557 |
| Dye | BioTracker NucView 405 Blue Caspase-3 Dye | Merck KGaA | 23N0609 |
| Software | RStudio | RStudio | 2022.07.1 Build 554 |
| Software | GraphPad Prism | GraphPad Software LLC | 11.0.0 |
| Software | R | R Project | 4.4.1 |
| Software | FlowJo | BD | v10.10.0 |
| Software | FACS Diva | BD | v8.0.1 |
| Software | CytExpert | Beckman Coulter | v2.4.0.28 |

**Supplementary Table 6: Materials and Software**
